# Neuronal activity patterns regulate BDNF expression in cortical neurons via synaptic connections and calcium signaling

**DOI:** 10.1101/2021.02.28.433239

**Authors:** Yumi Miyasaka, Nobuhiko Yamamoto

## Abstract

During development, cortical circuits are remodeled by spontaneous and sensory-evoked activity via alteration of the expression of wiring molecules. An intriguing question is how physiological neuronal activity modifies the expression of these molecules in developing cortical networks. Here, we addressed this issue, focusing on brain-derived neurotrophic factor (BDNF), one of the factors underlying cortical wiring. Real-time imaging of Bdnf promoter activity in organotypic slice cultures revealed that patterned stimuli differentially regulated the increase and the time course of the promoter activity in upper layer neurons. Calcium imaging further demonstrated that stimulus-dependent increases in the promoter activity were roughly proportional to the increase in intracellular Ca^2+^ concentration per unit time. Finally, optogenetic stimulation showed that the promoter activity was increased efficiently by patterned stimulation in defined cortical circuits. These results suggest that physiological stimulation patterns differentially tune activity-dependent gene expression in developing cortical neurons via cortical circuits, synaptic responses, and alteration of intracellular calcium signaling.

## Introduction

Neuronal activity plays a crucial role in the formation of functional connections during development. This activity-dependent circuit formation has been well characterized in the sensory cortex: Spontaneous and sensory-evoked neuronal activity remodels neuronal connections in the visual and somatosensory cortex (1–4). Moreover, such remodeling is regulated by activity-dependent expression of molecules that affect axon and dendrite behavior, indicating that neuronal activity is adequately converted into molecular signals (5–14).

The next challenge is to understand how physiological neuronal activity patterns affect the gene expression that regulates cortical wiring. Indeed, neuronal firing is known to occur with characteristic frequency *in vivo* during cortical development (15–17). For example, long-beta oscillation and gamma rhythmic activity are prominently generated spontaneously or in a stimuli-evoked manner during the early postnatal period. Theses characteristic neural activities are thought to contribute to circuit remodeling, but few studies have investigated how patterned activity influences gene expression (18, 19).

To address this issue, we focused on brain-derived neurotrophic factor (BDNF), one of the best characterized activity-dependent molecules, which is expressed in upper layer neurons (20–22) and affects cortical wiring by promoting axonal and dendritic growth (23–30). Transcriptional regulation of the Bdnf gene, which responds to neuronal activity, has also been studied biochemically (31–35). Fundamentally, Ca^2+^ influx triggers activation of calcium-dependent transcription factors such as cAMP response element binding protein (CREB), and their binding to specific DNA sites induces the expression of downstream genes (36–41). Such characterization allows us to study the activity-dependent gene regulation mechanism from the physiological and molecular biological points of view.

In the present study, we investigated how neuronal activity patterns modify Bdnf promoter activity in living individual cortical neurons. For real-time imaging of the promoter activity, a luciferase expression vector under the control of the Bdnf exon IV promoter, which is the best characterized promoter in the activity-dependent context, was electroporated into upper layer neurons in organotypic cortical cultures. In this culture, cortical cytoarchitecture is preserved, and intracortical connections such as horizontal connections that connect upper layer neurons are formed in an activity-dependent fashion, as they are *in vivo* (12, 42–46). BDNF promoter activity as well as calcium dynamics were then investigated in upper layer neurons by stimulating the cortical circuits electrophysiologically and optogenetically with various stimulus patterns. The results demonstrated that patterned neuronal activity differentially modulates Bdnf expression via alteration of the calcium signaling within the cortical networks.

## Results

### Bdnf promoter activity changes temporally in individual cortical cells in response to chemical stimulation

To study neuronal activity-dependent Bdnf promoter activity in individual cortical neurons, we performed live imaging with a luciferase assay in organotypic rat cortical cultures (42). The promoter activity was investigated in upper layer neurons, as they have BDNF-dependent plasticity in terms of development and circuit formation (24, 27, 47, 48). A luciferase expression plasmid containing Bdnf promoter IV (Bdnf-luc plasmid), which primarily underlies activity-dependent Bdnf expression (38, 49), was used for live imaging (Fig. 1A). The promoter region is located immediately upstream of the transcription initiation site and is highly conserved among species (Fig. S1) (50). The Bdnf-luc plasmid was electroporated sparsely into upper layer cells (Fig. 1A) (12).

**Fig. 1.**
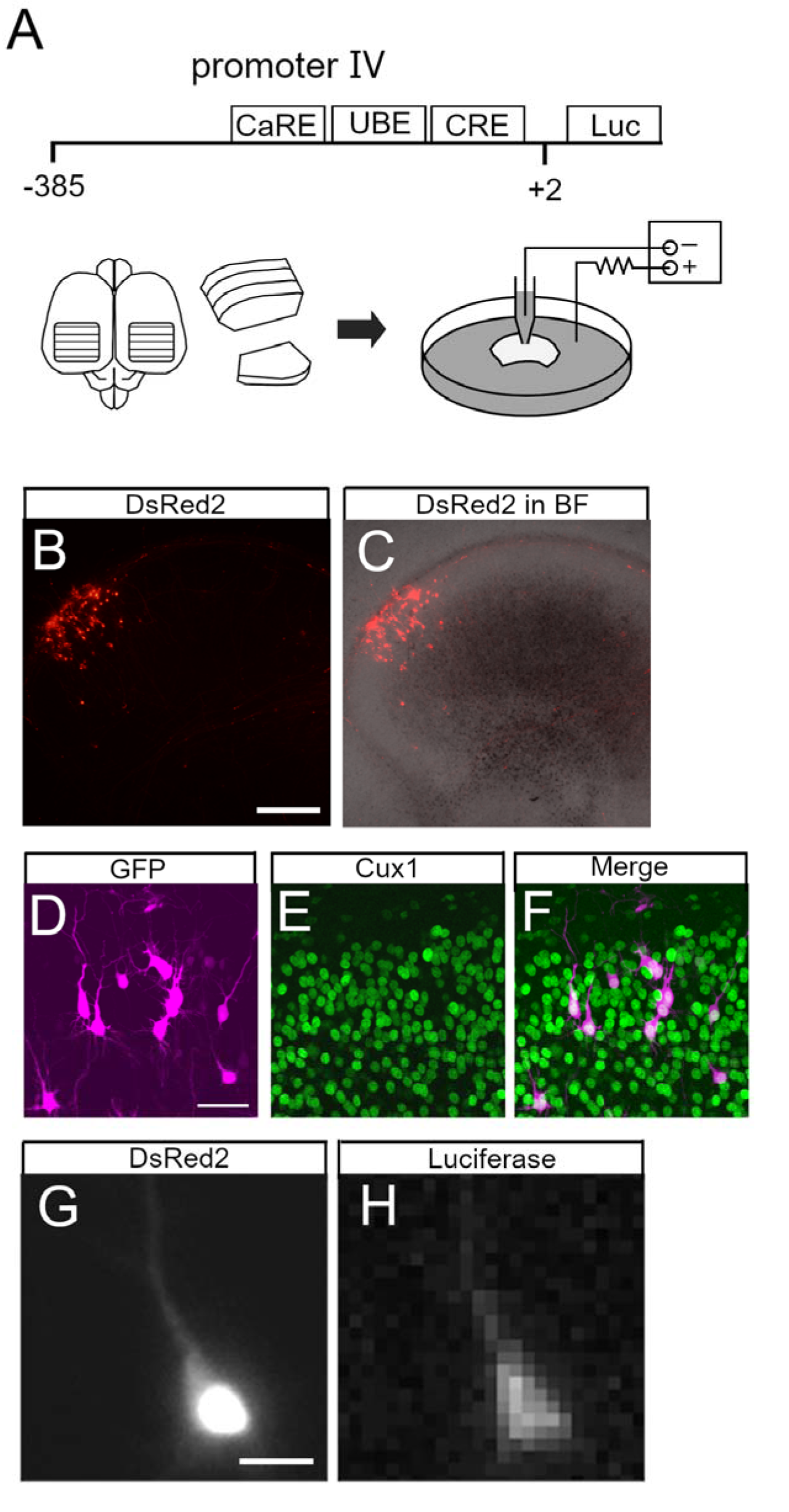
Live imaging of Bdnf promoter activity in upper layer neurons. **A**, Bdnf promoter region and schematic representation of organotypic cortical slice culture and electroporation. CaRE, calcium response element; UBE, upstream stimulatory factor binding element; CRE, cAMP-response element; Luc, luciferase. **B, C**, Electroporated (DsRed2-labeled) cells in a cortical slice. Fluorescence image (**B**) and merged image with bright field (BF) (**C**). **D–F**, Cux1 immunoreactivity in the electroporated cells. Electroporated neurons (**D**) with immunostaining for Cux1 (**E**) and merged image (**F**). **G, H**, Luciferase signal in an electroporated neuron. The neuron exhibits DsRed2 (**G**) and luciferase signals (**H**). Scale bars: B, 500 μm; D, 50 μm; G, 20 μm.

Enhanced green fluorescent protein (EGFP)- or DsRed2-encoding plasmid was co-transfected to identify electroporated cells readily and to reveal cellular morphology. As shown in Fig. 1B and C, fluorescent protein-labeled cells were distributed in the upper layers (0.1 to 0.5 mm from the pial surface) of cortical explants.

Immunohistochemistry after live imaging showed that the transfected cells were mostly Cux1-immunopositive (Fig. 1D–F). Roughly half of the fluorescent protein-labeled cells exhibited bioluminescent signals in the presence of luciferin (Fig. 1G and H).

We then investigated temporal changes of Bdnf promoter activity using KCl treatment (final concentration, 25 mM) at one week *in vitro,* when spontaneous neuronal activity is very low (12). Endogenous Bdnf expression in cortical explants was confirmed to increase markedly upon treatment (Fig. S2A). For quantification of the promoter activity in individual cells, the bioluminescent signals were normalized by the baseline intensity in each cell and are referred to here as “luciferase signals” (see Materials and Methods). In the absence of KCl, the luciferase signals in individual cells were almost unchanged for up to 20 h (Fig. 2A and D), but the signals in most cells started to increase within 2 h after the initiation of KCl treatment, and reached a maximal value after approximately 10 h (Fig. 2B and E, movie1). To characterize the increase in Bdnf promoter activity, we measured the peak amplitude, total signal, and slope of the luciferase signals, which reflect the maximum level, gross amount, and rapidity of Bdnf expression, respectively (see Materials and Methods). These parameters were considerably larger in KCl-treated than in untreated cultures (6.1 ± 0.28*** vs 1.2 ± 0.027 for peak amplitude, 65 ± 3.5*** vs -1.6 ± 0.83 for total signal, 0.92*** ± 0.046 vs 0.098 ± 0.017 for slope, *n* = 123 cells from ten KCl-treated cultures and *n* = 22 cells from two untreated cultures, \*\*\**P* < 0.0001, Mann–Whitney *U* test) (Fig. 2G–I). Interestingly, the increase in Bdnf promoter activity varied widely among cells (Fig. 2E and G). The peak amplitude was not related to the baseline intensity (Fig. S3), suggesting that the luciferase signals in individual cells are not due to transfection efficiency, but reflect their own expression levels.

**Fig. 2.**
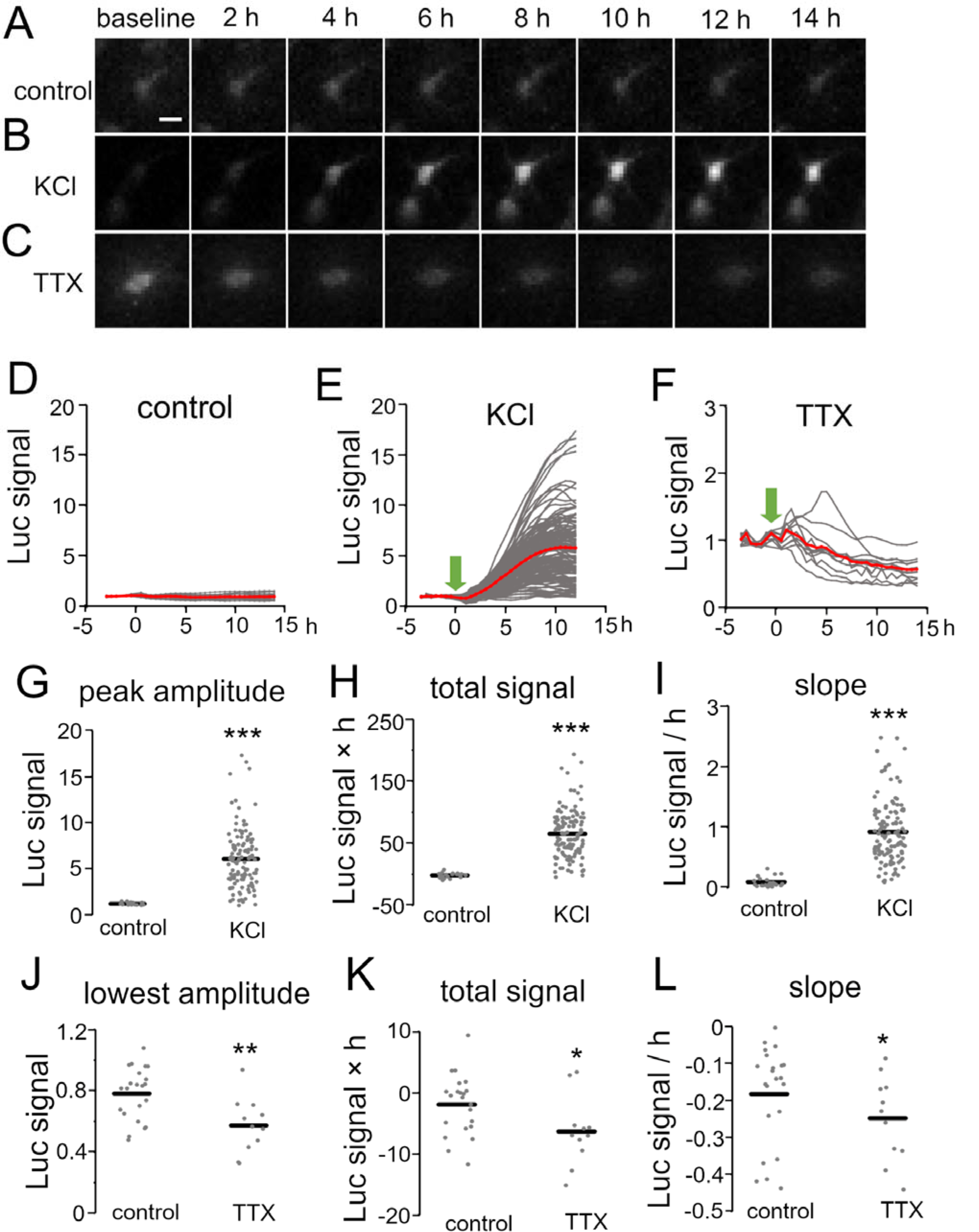
Temporal properties of Bdnf promoter activity in response to pharmacological stimulation. **A–C**, Time lapse images of luciferase signals without treatment (**A**) and with KCl (**B**) and TTX (**C**) treatment. Each picture was taken at the indicated time. Scale bar, 20 μm. **D–F**, Time courses of luciferase signals in individual neurons without treatment (**D**) and with KCl (**E**) and TTX (**F**) treatment. Gray lines indicate luciferase signals in each neuron, and red lines show averages. Arrows indicate the initiation of the treatment. **G–L**, Quantitative analysis of KCl (**G–I**) and TTX treatment (**J–L**). The peak amplitude (**G**), lowest amplitude (**J**), total signals (**H, K**), and slopes (**I, L**) were analyzed. Horizontal bars represent the average values. Asterisks indicate a significant difference compared to control (Mann–Whitney *U* test, \**P* < 0.05, ** *P* < 0.01, *** *P* < 0.001).

The effect of neuronal activity on Bdnf promoter activity was also examined by suppressing neuronal firing at 2 weeks *in vitro*, when spontaneous activity is prominent (see below) (12). To inhibit spontaneous firing activity, a sodium channel blocker, tetrodotoxin (TTX), was applied to the culture medium (final concentration, 100 nM).

In accordance with a considerable decrease in endogenous Bdnf expression (Fig. S2B), the luciferase signal gradually decreased and reached its lowest amplitude approximately 10 h after TTX addition (0.57 ± 0.054** vs 0.78 ± 0.036 for lowest amplitude, -6.3 ± 1.7* vs -1.9 ± 1.0 for total signal, -0.19 ± 0.025* vs -0.12 ± 0.018 for slope, *n* = 12 cells from two TTX-treated cultures, \*\**P* < 0.01, * *P* < 0.05, Mann– Whitney *U* test) (Fig. 2C, J–L). Thus, pharmacological treatments demonstrated that Bdnf promoter activity was temporally and differentially regulated in individual cortical neurons by neuronal activity.

As described above, Bdnf promoter activity differed considerably among cells (Fig. 2E). We investigated whether this individual variability is related to the spatial arrangement of cortical neurons. We found that cells with similar peak amplitude of the luciferase signal after KCl treatment tended to cluster (Fig. S4A). Quantitative analysis showed that the proportion of “similar pairs”, cell pairs with similar promoter activity, was significantly higher (\**P* < 0.05, \*\**P* < 0.01, Fisher’s Exact Test) in cell pairs that were located close to each other (intercellular distances < 30 μm) than in more widely separated cell pairs (intracellular distances > 30 μm) (Fig. S4B–D, Table S1, see Materials and Methods). These findings suggest that cortical neurons with similar activity dependence on Bdnf expression are located in close proximity to each other under the uniform depolarization of KCl treatment.

### Patterned stimuli differentially alter Bdnf promoter activity

Next, we investigated how patterned neuronal activity influences Bdnf promoter activity. As Bdnf expression in physiological conditions is regulated postsynaptically by excitatory synaptic inputs (20, 22, 51), the promoter activity was examined by stimulating inputs after 2 weeks in culture, when intrinsic cortical connections are established with synapse formation (12, 42, 52). To clarify the effects of the excitatory inputs, inhibitory transmission was suppressed by adding a GABAA receptor blocker, picrotoxin (PTX), to the culture medium. However, a complication was the occurrence of frequent spontaneous firing activity at this stage (Fig. S5) (12), which interferes with the assessment of the role of evoked activity on the promoter activity. To reduce the spontaneous activity without perturbing synaptic responses, the concentrations of Ca^2+^ and Mg^2+^ in the extracellular medium were raised (53). Under this condition, spontaneous firing activity was almost abolished (Fig. S5) without diminishing activity-dependent transcription (Fig. S6 and Table S2).

Conventional electrical stimulation was carried out with a pair of platinum wires that were embedded into culture dishes. These electrodes penetrated the cortical slice away from the Bdnf-luc-transfected region to avoid antidromic activation (Fig. S7A). Calcium imaging with GCaMP6f or Oregon Green 488 BAPTA-1 (OGB-1) showed that upper layer neurons faithfully responded to each stimulation (Fig. S7B, movie 2). These responses were almost abolished with DNQX and D-AP5 (Fig. S7B), indicating that evoked responses were largely due to excitatory synaptic activation.

It is known that characteristic neuronal activities are generated in the developing brain (15, 17, 54). To mimic the occurrence of such neuronal activities, we applied constant-frequency stimulation (0.1 and 2 Hz for delta band, 10 Hz for alpha band, 20 Hz for beta band, and 60 Hz for gamma band) and burst stimulation (60-Hz burst at 0.33 Hz for gamma burst and 100-Hz burst at 5 Hz for theta burst). Luciferase signals were then measured in individual upper layer neurons. We found that some patterned stimuli clearly increased the luciferase signals, but others seemed to be ineffective (Fig. 3A–C). Indeed, the time courses of the luciferase signals showed that Bdnf promoter activity increased differently by the stimulation patterns (Fig. 3E–M). Moreover, as was the case with KCl treatment, not all transfected cells exhibited positive responses (Fig. 3E–L). These responses were confirmed to be evoked synaptically, since the stimulation did not increase the luciferase signals in the presence of synaptic blockers (Fig. 3D).

**Fig. 3.**
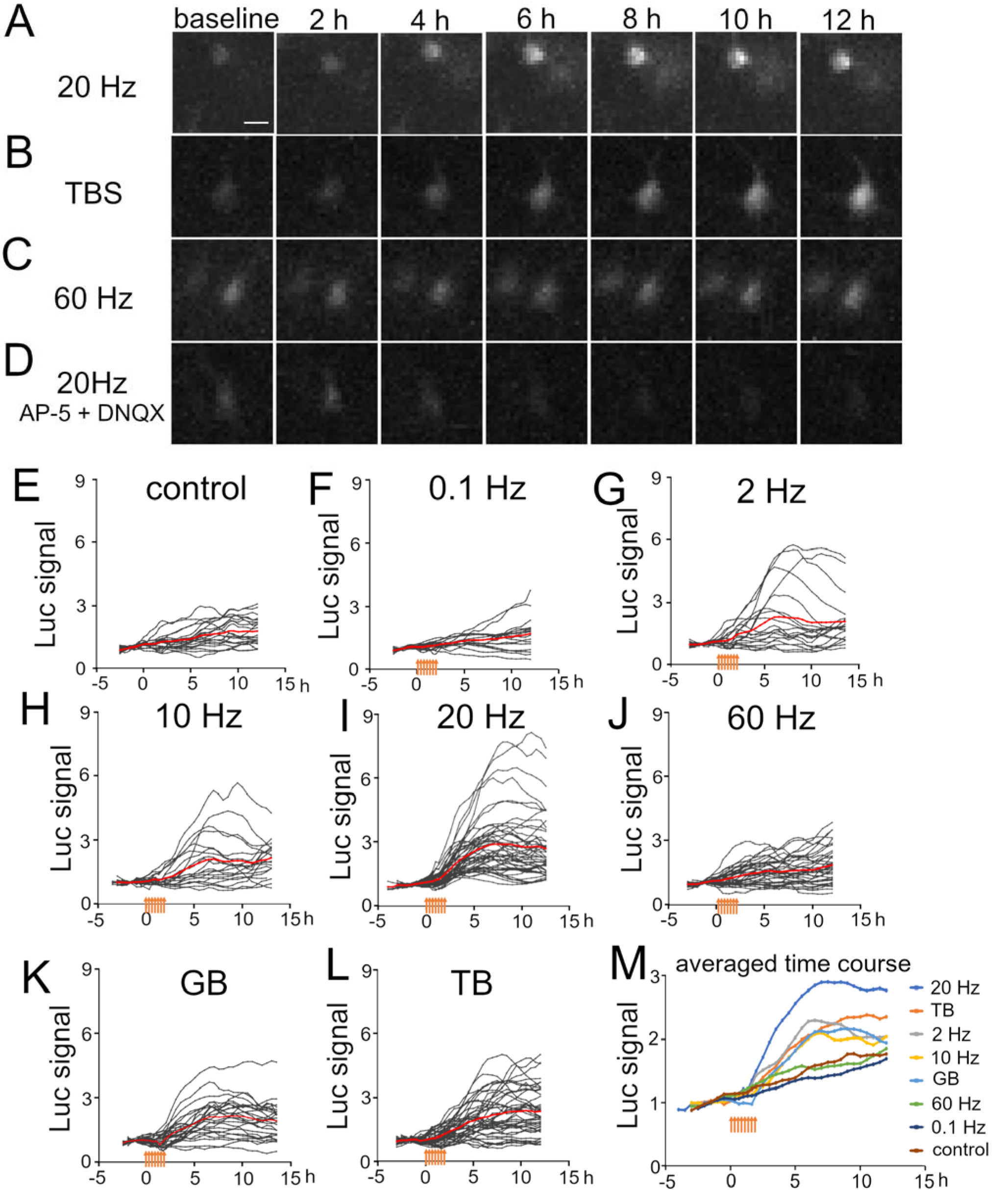
Temporal properties of Bdnf promoter activity in response to electrical stimulation at various frequencies. **A–D**, Time lapse images of luciferase signals for 20 Hz stimulation (**A**), TB stimulation (**B**), 60 Hz stimulation (**C**), and 20 Hz stimulation in the presence of AP-5 and DNQX (**D**). Each picture was taken at the indicated time. Scale bar, 20 μm. **E–L**, Time courses of luciferase signals in individual neurons for control (**E**), 0.1 Hz (**F**), 2 Hz (**G**), 10 Hz (**H**), 20 Hz (**I**), 60 Hz (**J**), GB (**K**), and TB (**L**) stimulation. Gray lines indicate luciferase signals in each neuron, and red lines show the averages. **L**, Averaged time courses of the luciferase signals for each stimulation. The signals were averaged from all cortical neurons shown in G–K. Orange arrows indicate stimulus time points.

Luciferase signals tended to increase slightly and gradually even without stimulation, probably due to removal of inhibitory transmission (Fig. 3E). To quantify the effects of each stimulation pattern, the total signals and slopes in Bdnf promoter activity were compared before and after stimulation (without stimulation total signal, 7.0 ± 1.4 vs 7.5 ± 1.2; slope, 0.18 ± 0.026 vs 0.15 ± 0.023, *n* = 21 cells from 5 cultures, Fig. 3E). We found that stimuli with 2 Hz, 10 Hz, 20 Hz, gamma burst (GB), and theta burst (TB) were effective in increasing the promoter activity (Fig. 4A and B). The total signals were 2- to 4-fold larger after applying these stimuli (after vs before, 14 ± 4.5 vs 4.8 ± 0.97 for 2 Hz, *n* = 19 cells from 3 cultures; 13 ± 3.1* vs 4.9 ± 2.4 for 10 Hz, *n* = 20 cells from 5 cultures; 23 ± 3.0*** vs 4.6 ± 0.63 for 20 Hz, *n* = 46 cells from 4 cultures; 16 ± 2.4*** vs 3.4 ± 1.3 for GB, *n* = 26 cells from 3 cultures, 17 ± 2.2*** vs 2.8 ± 1.1 for TB, *n* = 40 cells from 4 cultures, \*\*\**P* < 0.001, \**P* < 0.05, Mann–Whitney *U* test), and the slopes were 3- to 6-fold larger after 2 Hz, 10 Hz, 20 Hz, GB, and TB (after vs before, 0.35 ± 0.086* vs 0.097 ± 0.019 for 2 Hz, 0.31 ± 0.067*** vs 0.049 ± 0.024 for 10 Hz, 0.48 ± 0.048*** vs 0.092 ± 0.013 for 20 Hz, 0.35 ± 0.049*** vs 0.067 ± 0.027 for GB, 0.30 ± 0.041*** vs 0.056 ± 0.021 for TB, \*\*\**P* <0.001, \**P* <0.05, Mann–Whitney *U* test). In contrast, 0.1 Hz and 60 Hz stimulation elevated neither the total signals (after vs before, 6.9 ± 2.0 vs 5.6 ± 0.86 for 0.1 Hz, *n* = 16 cells from 2 cultures, 6.8 ± 1.1 vs 6.2 ± 0.91 for 60 Hz, *n* = 37 cells from 3 cultures, *P* > 0.05, Mann–Whitney *U* test) nor the slopes (after vs before; 0.13 ± 0.027 vs 0.11 ± 0.017 for 0.1 Hz, 0.14 ± 0.019 vs 0.12 ± 0.018 for 60 Hz, *P* > 0.05, Mann–Whitney *U* test) (Fig. 4A and B). Next, the proportion of cells that responded to the electrical stimulation was examined. Stimulation with 20 Hz, GB, and TB activated roughly 70% of the analyzed cells (36/46 for 20 Hz, 18/26 for GB, 28/40 for TB), but the other stimulation patterns activated only around 20% of the cells (2/16 for 0.1 Hz, 6/19 for 2 Hz, 6/20 for 10 Hz, 9/37 for 60 Hz) (Fig. 4C). The multiplication of the total signal and the activated cell proportion were further calculated to estimate the total effectiveness. As shown in Fig 4D, 20 Hz stimulation was the most effective, followed by TB and GB stimulation. These results indicate that different patterned activity differentially regulated the transcriptional activity.

**Fig. 4.**
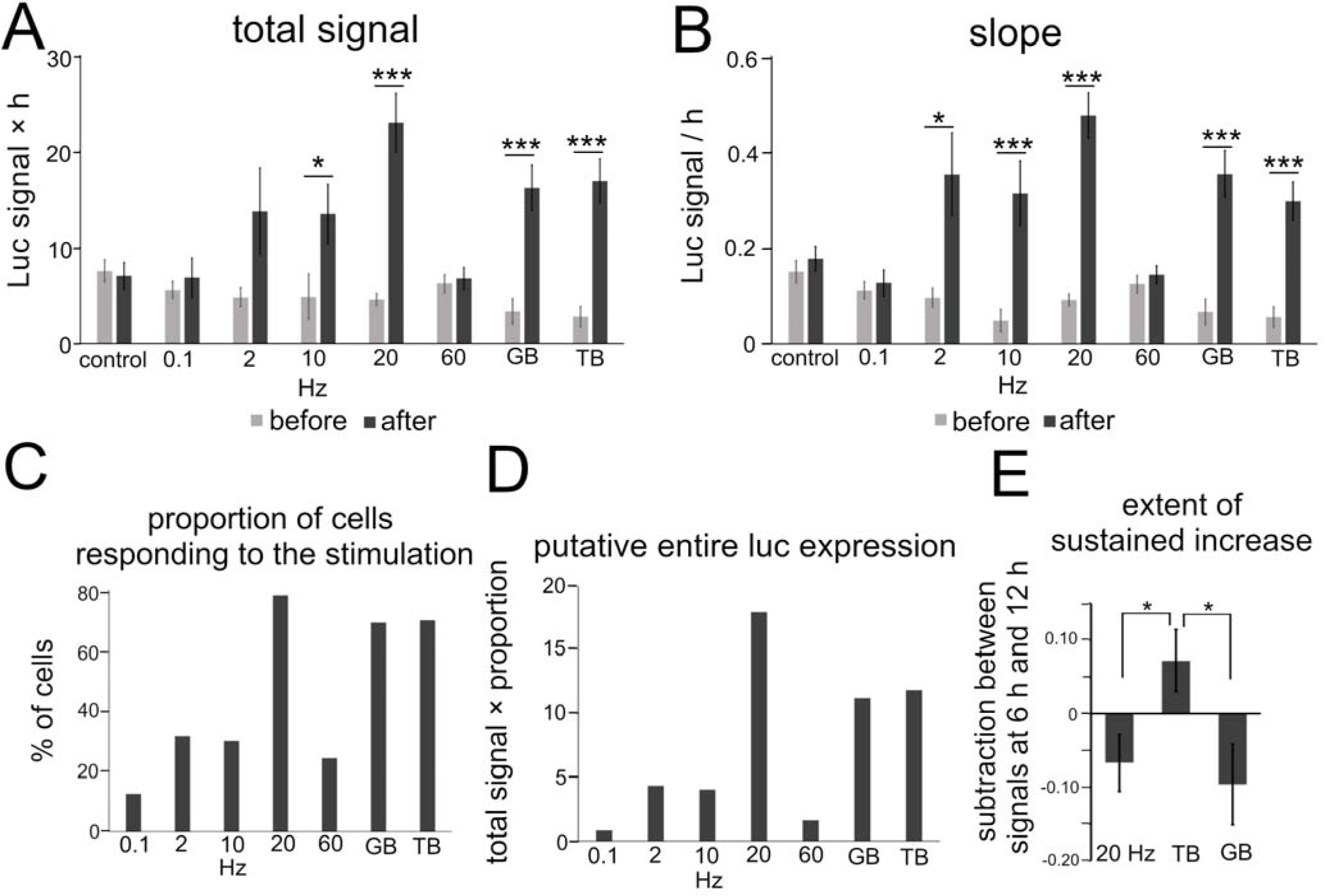
Quantitative analysis of the effects of patterned electrical stimulation. **A, B**, Increased levels for total signals (**A**) and slope (**B**). **C**, Proportion of cells responding to each stimulation. **D**, Putative entire luciferase expression. **E**, Extent of the sustained increase for 20 Hz, GB, and TB stimulation. Asterisks indicate a significant difference between the signals before and after stimulation (**A, B**) or among 20 Hz, GB, and TB stimulation (**E**) (Mann–Whitney *U* test or Tukey’s multiple comparison test, \**P* < 0.05, ** *P* < 0.01, *** *P* < 0.001). Bars represent the mean ± SEM.

Regarding the time course, the luciferase signals started to increase in every stimulation pattern within 2 h after the initiation of stimulation, but the profiles of time courses varied depending on the stimulation pattern (Fig. 3M). We analyzed the time courses of luciferase signals in response to 20 Hz, TB, and GB stimulation, which were most effective in inducing the promoter activity (Fig. 4D). In the cases of 20 Hz and GB stimulation, the signals mostly peaked around 6 h after stimulation, and then plateaued or slightly decreased (Fig. 3I, K, M). In contrast, TB stimulation tended to increase the signals in a gradual fashion until the end of the experiment session (Fig. 3L and M).

Quantitative analysis of the time courses showed that the extent of the sustained increase was significantly larger (-0.067 ± 0.039 for 20 Hz, -0.096 ± 0.055 for GB, 0.072 ± 0.043* for TB, Tukey’s multiple comparison test, \**P* < 0.05) in TB stimulation (Fig. 4E), suggesting that the time course of Bdnf promoter activity is also modulated by the stimulation patterns.

### Stimulus-dependent increase in Bdnf promoter activity is related to transient changes of intracellular Ca^2+^ concentration

To investigate how patterned stimuli differentially regulated Bdnf promoter activity, we investigated intracellular Ca^2+^ levels in each stimulation pattern, because previous studies have shown that Bdnf promoter activity is regulated by calcium-dependent mechanisms (33). Calcium imaging with OGB-1 showed that the intracellular Ca^2+^ concentrations were significantly suppressed during 20 Hz stimulation in the presence of nifedipine and APV, blockers of L-type voltage-gated Ca^2+^ channel and NMDA channel, which are the main source of Ca^2+^ influx (approximately 50% reduction, *n* = 3 cultures, Fig. 5A and B). In this condition, 20 Hz stimulation, which is the most effective to increase the promoter activity, did not increase the total signal or the slope after stimulation (total signal, 7.3 ± 1.5 vs 12 ± 2.9; slope, 0.19 ± 0.029 vs 0.25 ± 0.059, n = 11 cells from 2 cultures, *P* > 0.05, Mann–Whitney *U* test) (Fig. 5C–E).

**Fig. 5.**
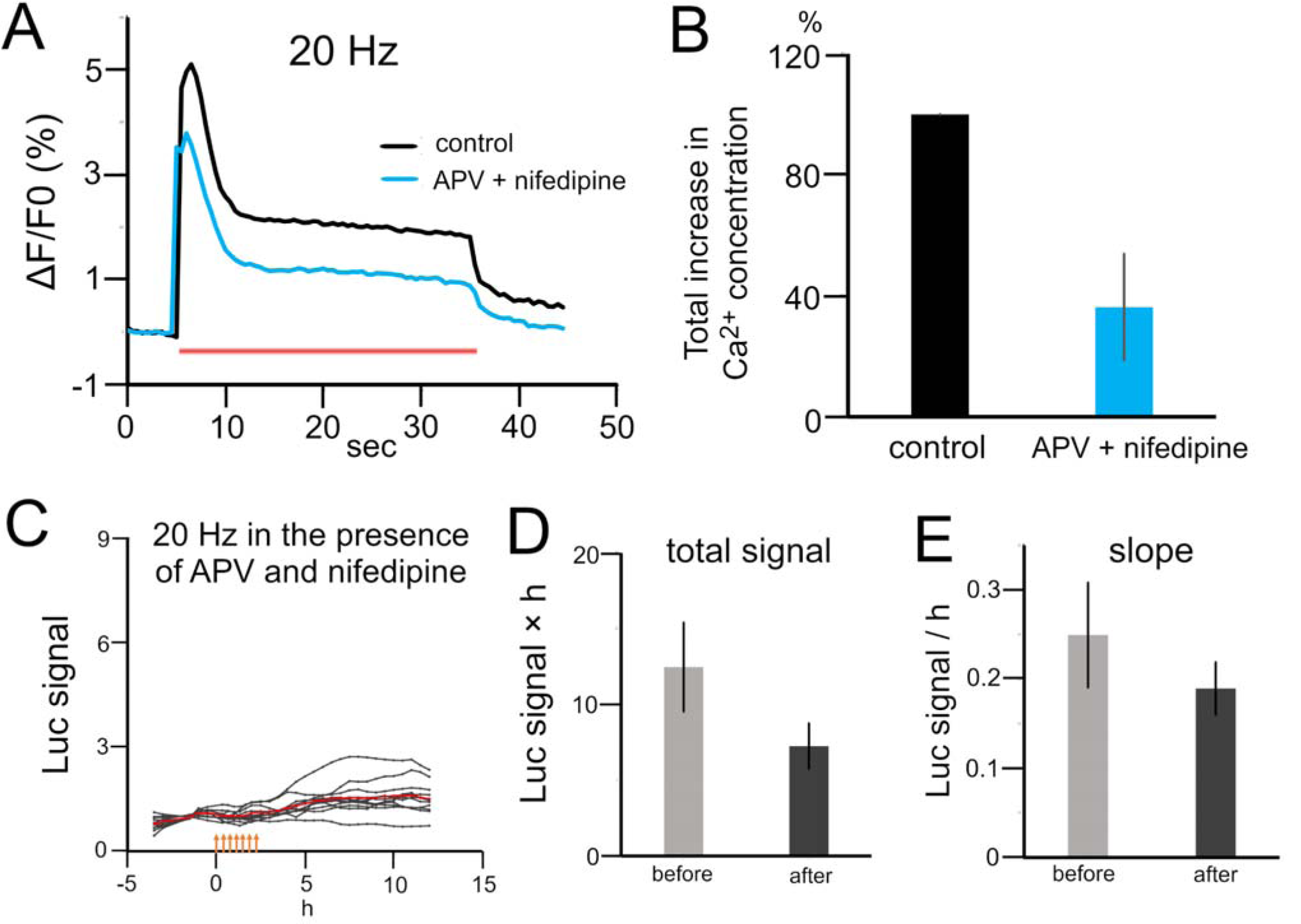
Effects of calcium channel blockers on stimulation-induced luciferase expression. **A**, Calcium signaling during 20 Hz stimulation in the presence or absence of APV and nifedipine. A red bar represents the period of stimulation. **B**, Quantitative analysis of the intracellular Ca^2+^ concentration with and without APV and nifedipine (n=3). **C**, Time courses of luciferase signals with 20 Hz stimulation in the presence of APV and nifedipine. Orange arrows indicate stimulus time points. **D, E**, Changes in the total signal (**D**) and slope (**E**) in the presence of APV and nifedipine.

This result confirms that electrical stimulation up-regulated Bdnf promoter activity via intracellular calcium signaling.

We hypothesized that the different increases in Bdnf promoter activity in response to stimulation patterns are due to the distinct intracellular Ca^2+^ concentrations. To test this, the calcium responses to each patterned stimulation were measured.

Calcium imaging showed that different stimulation patterns evoked different calcium oscillations (Fig. 6A–F). The increased Ca^2+^ concentrations and those per unit time were also different among the stimulation patterns. The total increase levels in Ca^2+^ concentrations (normalized by the maximal value) were 0.50 ± 0.12 for 0.1 Hz, 0.99 ± 0.0082 for 2 Hz, 0.18 ± 0.049 for 10 Hz, 0.13 ± 0.015 for 20 Hz, 0.022 ± 0.0016 for 60 Hz, 0.72 ± 0.0092 for GB, and 0.14 ± 0.012 for TB (*n* = 6 cultures, Fig. 6G).The increase levels per unit time (normalized by the maximal value) were 0.23 ± 0.053 for 0.1 Hz, 0.69 ± 0.047 for 2 Hz, 0.91 ± 0.037 for 10 Hz, 0.85 ± 0.044 for 20 Hz, 0.49 ± 0.041 for 60 Hz, 0.49 ± 0.054 for GB, and 0.96 ± 0.030 for TB (Fig. 6H). These values were plotted against the promoter activity obtained by the luciferase assay for each stimulation (Fig. 6I and J). The total luciferase signals were highly correlated (r = 0.69, correlation coefficient) with intracellular Ca^2+^ concentrations per unit time (Fig. 6J), whereas little correlation was found (r = -0.04) between the total luciferase signals and the total calcium levels (Fig. 6I). Thus, it is likely that physiological patterned activities differentially regulate Bdnf promoter activity through altering intracellular Ca^2+^ concentrations per unit time rather than the total calcium levels.

**Fig. 6.**
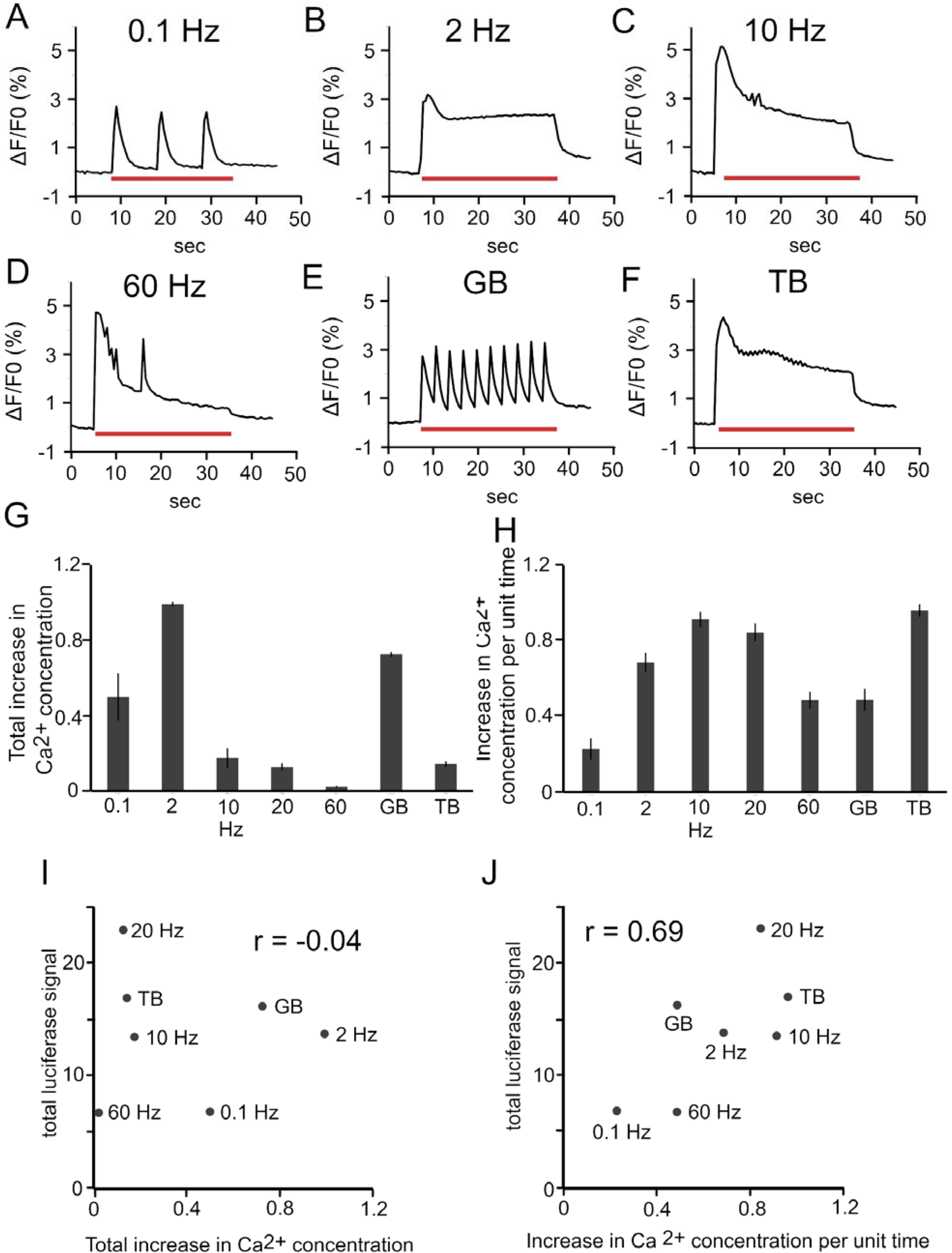
Relationship between intracellular Ca^2+^ concentration and luciferase signals. **A– F**, Changes in intracellular Ca^2+^ concentration (ΔF/F0) in upper layers during various stimulations. Red bars represent the period of stimulation. **G**, Total increases in intracellular Ca^2+^ concentration for each stimulation pattern (n = 6). **H**, Increases in intracellular Ca^2+^ concentration per unit time for each stimulation pattern. In (**G**) and (**H**), the value for each stimulation was calculated as the ratio to the maximum among the increases for all stimulation patterns, and averaged for the number of samples. **I**, Relationship between total luciferase signals and total changes in intracellular Ca^2+^ concentration. **J**, Relationship between total luciferase signals and changes in intracellular Ca^2+^ concentration per unit time.

### Bdnf promoter activity is induced in cortical neurons postsynaptically via upper layer neuronal circuits

Finally, we examined whether patterned neuronal activity regulates Bdnf promoter activity through physiologically and anatomically defined cortical circuits, focusing on intracortical connections between upper layer neurons (55, 56). For this, optogenetic stimulation was performed to stimulate upper layer neurons specifically. To stimulate a large number of neurons, a channel rhodopsin 2 (chr2) plasmid was transfected at E15 into mouse cortical neurons destined to become upper layer neurons using *in utero* electroporation prior to culturing. Cortical slices containing ChR2-expressing cells were then dissected, and the Bdnf-luc plasmid was transfected into the cells around ChR2-expressing cells (Fig. 7A). In this condition, upper layer neurons including the Bdnf-luc-expressing cells were surrounded by ChR2-expressing fibers (Fig. S8). Luciferase signals were then observed by applying blue light photostimulation. This optogenetic stimulation should produce only excitatory synaptic responses in luciferase-positive cells, since the chr2 plasmid is transfected only into excitatory neurons by the *in utero* electroporation technique.

**Figure 7.**
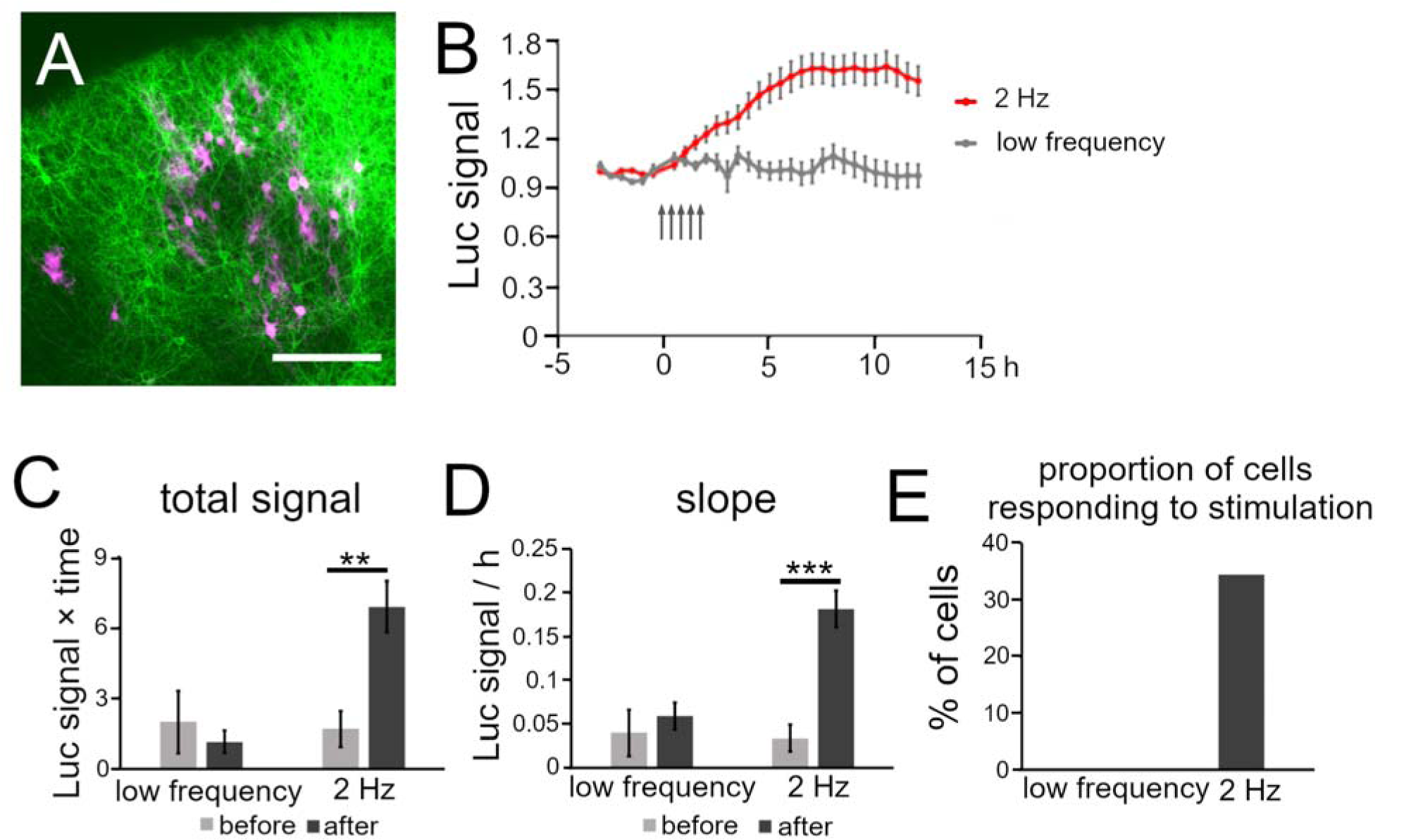
Analysis of Bdnf promoter activity in response to optogenetic stimulation. **A**, Representative confocal image of a cortical slice electroporated with ChR2 and luciferase vectors (green, ChR2-positive and red, luciferase-positive cells). Scale bar, 200 μm. **B**, Time courses of luciferase signals in each stimulation pattern. Arrows indicate the timing of stimulation. Bars represent SEM. **C, D**, Quantitative analysis of the increased level of luciferase signals for total signals (**C**) and slope (**D**). Bars represent SEM. Asterisks indicate a significant difference between before and after stimulation (Mann–Whitney *U* test, ** *P* < 0.01, *** *P* < 0.001). **E**, Proportion of cells responding to each stimulation.

Whether the Bdnf promoter was activated similarly in mouse cortical neurons was confirmed prior to the optogenetic stimulation experiment. Pharmacological treatment showed that the promoter activity was remarkably increased by KCl treatment (Fig. S9A). The increased levels of Bdnf promoter activity were comparable with those in rat neurons (Fig. S9B–D, Table S2). The increases and temporal properties of Bdnf promoter activity after patterned stimulation were also similar to those for rat neurons (Fig. S10, Table S3). Thus, neuronal activity modulates the promoter activity in a similar manner in both mice and rats.

Photostimulation with less than 2 Hz was applied to the cortical slices, as ChR2 hardly allows stable activation at high frequencies (57–59). Within 2 h after 2 Hz stimulation luciferase signals started to increase, and reached a peak at around 6 h and plateaued (Fig. 7B). The time course was similar to that elicited by electrical stimulation. Quantitative analysis showed that both the total signal and slope after 2 Hz stimulation were significantly larger than before stimulation (total signal, 6.9 ± 1.1** vs 1.7 ± 0.75; slope, 0.18 ± 0.021** vs 0.034 ± 0.015, *n* = 32 cells from 4 cultures, \*\**P* < 0.01, Mann– Whitney *U* test) (Fig. 7C and D). Moreover, this optogenetic stimulation activated a certain population (11 out of 32 cells) of cortical cells (Fig. 7E). In contrast, lower frequencies of stimulation (0.1 and 1 Hz) did not efficiently induce the total signals or slopes (total signal, 1.2 ± 0.46 vs 2.0 ± 1.3; slope; 0.059 ± 0.016 vs 0.040 ± 0.026, *n* = 11 cells from 2 cultures, *P* > 0.1, Mann–Whitney *U* test), and activated none of the cortical cells tested (0 out of 11 cells) (Fig. 7C–E). This result indicates that patterned excitatory synaptic inputs are responsible for the modulation of Bdnf promoter activity via a defined cortical circuit.

## Discussion

In the present study, we demonstrated spatiotemporal regulation of Bdnf promoter activity in the cortex by pharmacological and physiological stimulation. Our results show that patterned stimuli differentially upregulate Bdnf promoter activity in individual upper layer neurons, via adequate increases of intracellular Ca^2+^ concentration. In addition, cortical cells showing similar activity dependence for gene expression are spatially localized. These results suggest that physiological stimuli differentially tune activity-dependent gene expression in individual cortical neurons via intracellular calcium signaling, which is controlled by intrinsic cortical cell properties and synaptic inputs.

### Modulation of Bdnf promoter activity by patterned stimulation

We demonstrated the precise time course and elevation of stimulus-induced Bdnf promoter activity in individual cortical cells. A remarkable aspect is that the levels of increase were modulated differentially by stimulus patterns. Indeed, distinct patterned activities have been found in the developing cortex, and they are thought to contribute to circuit refinement (17). Here, we showed that patterned stimulation such as GB, TB, and 20 Hz (long beta-oscillation) stimulation efficiently induced Bdnf expression, although high delta (2 Hz) and alpha waves (10 Hz) were only moderate inducers.

These results suggest that neuronal activities, which are generated in the physiological situation, regulate molecular expression, which leads to the establishment of cortical wiring. In accordance with this view, recent studies have shown that patterned firing activity regulates guidance molecule expression (9, 13). Moreover, the present optogenetic experiment demonstrated that patterned activation of pre-synaptic fibers increased Bdnf promoter activity, suggesting that physiological neuronal activity modulates the gene expression in individual cortical neurons via defined connections.

How do patterned activities differentially regulate the increase of Bdnf expression? The present results not only confirmed the necessity of the increase in intracellular Ca^2+^ concentration but also indicated that a phasic increase is crucial for downstream molecular signaling. Ca^2+^-dependent transcriptional activity could be influenced by the temporally controlled Ca^2+^ concentration, which is differentially raised by stimulation patterns. Indeed, autophosphorylation of CaM kinase, which is critical for binding of transcription factors to Bdnf promoter IV, is regulated by Ca^2+^ stimulation at specific frequencies (60). The translocation of CREB-regulated transcription co-activator 1 (CRTC1) to the nucleus was also promoted by patterned stimulation (61). It is likely that specific patterned neuronal firing, regardless of spontaneous or evoked activity, efficiently promotes Bdnf expression by increasing transient intracellular Ca^2+^ concentration in cortical cells and by upregulating the transcription together with cofactors. However, factors other than temporal changes in intracellular Ca^2+^ concentration may also be required for the upregulation of Bdnf expression, since temporal Ca^2+^ concentration changes and Bdnf expression were not completely proportional (Fig. 6I).

Furthermore, we also found that patterned activity modulated the time courses of Bdnf promoter activity: 20 Hz and GB stimulation increased the promoter activity and generated a peak, but TB stimulation elicited gradual and continuous increases.

Patterned stimulation may alter the temporal response by affecting the dynamics of transcription factors or their DNA binding. For instance, transcriptional activity of NF-κB, which also regulates Bdnf exon IV transcription, is modulated by the frequency of Ca^2+^ oscillation (62, 63). Considering that NF-κB is translocated into the nucleus in response to Ca^2+^ stimulation, patterned stimulation may differentially regulate the temporal mobility of the transcription factor.

### Responsiveness of individual cortical neurons

We also found different responsiveness in individual cortical neurons. The level of increase in promoter activity varied from cell to cell, even in uniform activation by KCl treatment, suggesting that activity-dependent responsiveness is intrinsically variable in cortical neurons. In support of this view, diversity in activity-dependent promoter activity has also been reported in dissociated cortical cells from Bdnf-luc transgenic mice (64). This may be due to different expression levels in calcium channels, cytoplasmic signaling molecules, and/or different phosphorylation levels of transcription factors, among cells (65, 66). These intrinsic biochemical properties may also contribute to the diversity of the proportion of responding cells in response to the stimulation patterns (Fig. 4C). In fact, the Bdnf exon IV promoter contains three critical domains for calcium-dependent transcription, which is activated by calcium pathways that initiate from distinct calcium channels and calcium-stimulated kinases (51).

Considering that the various kinds of calcium channels may be differently expressed in each cortical neuron (65), neuronal activities probably modify the proportion of responding cells by switching among types of activated calcium channels in response to activity patterns.

Regarding spatial distribution, cells with similar promoter activity were closely localized (Fig. S4), suggesting that cells with similar biochemical properties in their cytoplasmic molecular pathways tend to cluster. This spatial localization is in a very narrow range (0–30 μm), which is more restricted than the so-called columnar structures in the visual and somatosensory cortex (67, 68); rather, the localization is similar to the size of microcolumns (69, 70). Indeed, as cortical neurons originating from the same lineage have similar physiological properties and are spatially localized (71), these cells may have genetically similar properties. Moreover, this spatial profile was retained even by receiving effective physiological activities (Fig. S11 and Table S4), which suggests that spatial arrangement is unaffected by neuronal network activation.

In summary, patterned stimulation differentially modulates the level of increase and time course of Bdnf promoter activity via phasic changes in intracellular Ca^2+^ concentration and intrinsic biochemical properties within physiological neuronal circuits. Since BDNF released from cells promotes neurite outgrowth and branching of neighboring neurons (24, 25, 27), patterned neuronal activity may contribute to determination of the magnitude and time window in remodeling processes in neuronal wiring, and control local circuit formation by regulating Bdnf expression spatiotemporally.

### Materials and Methods Animals and ethics statement

All experiments were performed according to the guidelines established by the animal welfare committees of Osaka University and the Japan Neuroscience Society.

Sprague–Dawley (SD) rats (Nihon–Dobutsu) and ICR mice (Japan SLC and CLEA Japan) were used in this study.

### Organotypic slice culture

Organotypic cortical slice cultures were prepared as described previously (42). In brief, cortical slices were dissected from postnatal day (P) 1 rats or mice. The slices were placed on a membrane insertion (Millicell-CM; catalog no. PICMORG50; Millipore) coated with rat-tail collagen. The culture medium consisted of a 1:1 mixture of DMEM and Ham’s F-12 (Invitrogen) with several supplements (Yamamoto et al., 1992). The cultures were maintained at 37°C in an environment of humidified 95% air and 5% CO_2_.

### Construction of luciferase expression vector

Based on the sequence information of rat Bdnf (NC_005102), the exon4 promoter region (387 bp, from -384 to +2) was amplified by PCR with a pair of primers (5≠-GAATCCAGGTAGACAGCTTGGCAG-3≠ and 5≠-ACTGGGAGATTTCATGCTAGCTCG-3≠). The template genome was obtained from rat tail. The PCR product was cloned into pGEM-T-Easy vector (Promega), and the promoter region was further amplified by PCR using primer pairs containing recognition sites of restriction enzymes (5≠-TCTAGCTAGCAAAGAAAGAAAGAAAAAAGAAAAG-3≠ and 5≠-ACTACTCGAGGTGGGAGTCCACGAGAG-3≠). The PCR product was digested with the enzymes, and cloned into pGL3-basic vector (Promega), predigested with the same restriction enzymes.

### Local cell electroporation

To transfect the luciferase construct into upper layer cortical cells, electroporation with a glass microelectrode was performed after 2 days *in vitro* (DIV) as previously described (12). In brief, a mixture (0.5 μl) of Bdnf-luc plasmid (1.6 mg/ml) and pCAGGS-DsRed2 or pCAGGS-EGFP (0.4 mg/ml) in Hanks’ solution was applied to the surface of slices. Immediately afterwards, electrical pulses (10 trains of 200 square pulses of 1-msec duration, 200 Hz, 450 mA) were delivered with a glass microelectrode (300 µm diameter). For calcium imaging, CAG-GCaMP6f vector (2.0 mg/ml) (Addgene) was transfected into cortical cells in the same way.

### Quantitative reverse transcription-PCR analysis

After 1 or 2 weeks in culture, mRNAs were extracted from cortical explants that had been treated with KCl and TTX. After cDNA synthesis (Transcriptor First Strand cDNA Kit, Roche), gene expression was quantified by TaqMan Gene Expression Assay (Applied Biosystems). Rat GAPD was used as an endogenous control to normalize gene expression (Applied Biosystems). To amplify a specific sequence of Bdnf, a primer pair (5≠-AGCGCGAATGTGTTAGTGGT-3≠ and 5≠-GCAATTGTTTGCCTCTTTTTCT-3′) and a universal probe were used.

### Real-time luciferase assay

One day before the experiment, D-luciferin potassium salt (Wako) was added to the culture medium (final concentration, 0.1 mM). On the day of the experiment, cultured cortical slices were placed in an incubation chamber (Tokai Hit UK-A16U, 35°C, 5% CO_2_) on a microscope stage. After identifying DsRed2- or EGFP-positive cells, the luciferase signals were captured by an EMCCD camera (Andor, iXon3) attached to an upright microscope (Nikon, FN-S2N) through a 20x objective lens (NA, 0.5). The signal to noise ratio was increased by 4×4 binning, gain 1000, and 5- or 10- min exposure.

After confirming that the luciferase signals were stable on the microscope stage, the recording was started. The signals were recorded for 3 h before pharmacological, electrophysiological and optogenetic stimulation, and the slices were further observed for more than 12 h. The luciferase signals were taken at 30-min intervals.

### Pharmacological treatments

To increase neuronal activity, a high potassium solution (final concentration, 25 mM) was applied to the culture medium approximately 3 h after starting live imaging. To suppress neuronal firing, TTX (final concentration, 100 nM; Seikagaku-Kogyo) was applied to the medium. To block synaptic transmission, D-AP5 (100 µM) (Tocris) and DNQX (20 µM) (Tocris) were applied prior to imaging. To suppress spontaneous activity, the concentrations of Mg^2+^ and Ca^2+^ were raised to 4 mM (53). Picrotoxin (final concentration, 100 µM; nacalai tesque) was also applied to suppress the effect of inhibitory neurons.

### Electrophysiological stimulation

To apply electrical stimulation to cortical cells, we constructed hand-made electrode dishes, which are composed of a pair of platinum electrodes embedded in the bottom of culture dishes. One day before the experiment, the electrodes were inserted into cortical slices away from the pGL3-bdnf vector-transfected cells. For applying patterned activity, the electrodes were connected to an isolator (Bak Electronics, BSI-2), which was controlled by a programmable stimulator (A.M.P.I, Master-8). Various stimulation pulses (0.1 Hz for 20 min, 2 Hz for 10 min, 10 Hz for 2 min, 20 Hz for 1 min, 60 Hz for 20 s, GB for 10 min, and TB for 1 min) were then delivered to the slices (amplitude, 300 µA; duration 1 ms). GB and TB consist of a 60-Hz burst for 100 ms at 0.33 Hz and a 100-Hz burst for 30 ms at 5 Hz, respectively. A total of 1200 pulses were applied for each frequency, and the chunk of electrical pulses was repeated every 20 min up to 7 times.

### *In utero* electroporation

*In utero* electroporation was performed to express CAGGS-hChR2(H134R)-EYFP (ChR2-EYFP) in a large number of upper layer neurons. Pregnant mice at E15 were deeply anesthetized with isoflurane. The abdomen was surgically opened without opening the uterus itself. ChR2-EYFP (2 µg/µl) in PBS was injected into one cerebral ventricle. A tweezers-type platinum electrode was positioned beside the uterus, and square pulses (40 V; 50 ms) were delivered five times with electroporator (CUY20; BEX). After electroporation, embryos were allowed to develop until birth.

### Optogenetic stimulation

To apply stimulation to cortical cells, an optogenetic technique was also utilized. For this, a solid-state illuminator (475 nm peak wavelength; maximal power: 20 mW; Lumencor SPECTRA, Lumencor) was controlled by the programmable stimulator. The light intensity was 4.8 mW/mm^2^, which was enough to evoke the neuronal activity. Blue light stimulation (50 ms) was applied with various frequencies (0.1 – 2 Hz) through a 20x objective lens. The stimulation continued for 5 min and was repeated every 30 min up to 5 times.

### Immunohistochemistry

After the real-time luciferase assay, the slices were fixed for several hours with 4% paraformaldehyde in 0.1 M PBS. The slices were incubated at 4°C overnight with rabbit anti-Cux1 (1:250; Santa Cruz Biotechnology) and rat anti-GFP (1:1000; nacalai tesque). After extensive washes, the signals were visualized with Alexa 488-conjugated anti-rat IgG (1:500; invitrogen) and Cy5-conjugated anti-rabbit IgG (1:250; Jackson ImmunoResearch). The samples were embedded with 80% glycerol containing DAPI and DABCO, and observed by confocal microscopy through a 20× objective lens (Leica, TCS-SP5).

### Calcium imaging

Calcium imaging was carried out using GCaMP6f or Oregon Green 488 BAPTA-1 (Invitrogen) (OGB-1). A CAG-GCaMP6f vector (Addgene) was electroporated sparsely to cortical neurons in the slice culture at 2 DIV (see above). Alternatively, 4 µM OGB-1 including 0.01% Pluronic F-127 and 0.005% Cremophor EL were added to the culture medium for 30 min. The culture was washed with Hanks’ solution before imaging.

In both cases, calcium imaging was performed in the chamber on the microscope. Excitation light (475 µm wavelength) was applied to the observed area with the solid-state illuminator through 20× objective lens. Images were captured by an EMCCD camera at 4 Hz (exposure time 100 ms; binning 2×2; gain 300). To measure stimulus-induced calcium signals, the baseline (F0) was calculated by averaging the signals prior to stimulation, and stimulus-evoked changes (ΔF) divided by F0 were determined.

To evaluate the total increase levels for various stimulation patterns, stimulation with 0.1 to 60 Hz, GB, and TB burst was applied to the same cortical slices, and ΔF/F0 values were measured for a 0.4 x 0.4 mm area in the upper layers. The integral value of the ΔF/F0 values during the stimulation period was then calculated for each stimulation pattern, and was referred to as the total increase in the intracellular Ca^2+^ concentration. The total increase in the intracellular Ca^2+^ concentration was then divided by the stimulation period for each stimulation pattern, and was referred to as the total increase in the intracellular Ca^2+^ concentration per unit time. These parameters for each stimulation were divided by the maximum among the increases for all stimuli, and the ratios were compared between stimulus patterns.

### Recording of spontaneous activity

To examine spontaneous firing in cortical slices, extracellular recording was performed on multi-electrode dishes (interpolar distance, 0.3 mm), as described previously (12). In brief, the extracellular potentials were amplified and stored in a hard disk after digitization. Experiments were performed in the presence of 4 mM Mg^2+^ and Ca^2+^ and 100 μM picrotoxin.

### Image processing and analysis of temporal properties

Luciferase signals were analyzed with image processing software (ImageJ Fiji).

First, the region of interest was drawn within the cell body of each luciferase-positive cell, and the mean intensity was measured at every time point. The background signal outside the cell body was subtracted from the mean intensity. The subtracted value was smoothened by calculating the weighted moving average, and was defined as the signal intensity (L) at a given time point.

A baseline value (Lo) was calculated by averaging the signal intensities for 3 h prior to pharmacological or physiological stimulation. The ratio L/Lo was referred to as the “luciferase signal” to represent temporal changes of the luciferase signal.

For the pharmacological treatments, the peak and lowest amplitudes were defined as the maximum and minimum values after stimulation, respectively. The total signal was calculated by integrating the signals over time after stimulation. The slope was determined as the maximum inclination for 6 h after stimulation.

In the stimulation experiments, the luciferase signal gradually increased even in the baseline period, which may be due to picrotoxin in the culture medium. Therefore, the total signals and slopes were compared before and after stimulation to assess the effects of patterned activity on Bdnf promoter activity. The total signals after the stimulation were calculated by integrating the signals over time. The total signal before stimulation was calculated during the baseline period and extrapolated. The slopes were defined as described above. The percentage of responding cells was determined by calculating the percentage of cells exhibiting signals exceeding the threshold, which was 4 times the standard deviation of the signals before the stimulation. Putative entire luciferase expression was calculated by multiplying the total signals by the percentage of responding cells. The extent of sustained increase was calculated by subtracting the signal at 6 h from that at 12 h, and this value was normalized by the signal at 12 h.

### Analysis of spatial properties

For analysis of the spatial distribution of the cortical cells, the difference in the peak amplitude or the total signal between each cell pair was calculated. Cell pairs with a difference less than one-fourth of the maximal difference were defined as similar pairs, whereas those with a larger difference were defined as “different pairs”. Images of the analyzed neurons in fixed cortical explants were captured by confocal microscopy and the intercellular distance was measured by imageJ Fiji. The ratio of similar pairs was plotted in a histogram against the distance.

## Acknowledgements

This work was supported by Ministry of Education, Culture, Sports, Science and Technology KAKENHI on Dynamic Regulation of Brain Function by Scrap and Build System 16H06460 to N.Y., the Japan Society for the Promotion of Science KAKENHI Grants 15H04260 and 19H03325 to N.Y., and the Mitsubishi Foundation to N.Y. We thank Dr. Ian Smith for critical reading of the manuscript.

