## Supplemental materials for "Neuronal activity patterns regulate BDNF expression in cortical neurons via synaptic connections and calcium signaling"

Movies: 2

Supplementary tables: 4

Supplementary figures: 11

Movie 1. Live imaging of Bdnf promoter activity in response to depolarized stimulation. Time lapse imaging of luciferase activity in individual neurons before and after the depolarized stimulation. The movie shows changes of Bdnf promoter activity for 15 h before and after KCl application. “KCl” in red characters indicates the initiation time of stimulation. Scale bar, 50 μm.

Movie 2. Calcium responses of cortical neurons in response to electrical stimulation. Cortical neurons transfected with GCaMP6f were stimulated by platinum electrodes. “stim” in yellow characters indicates the stimulus point. Scale bar, 100 μm

Table S1. Distribution of cortical cells with similar promoter activity in response to depolarized stimulation

| Bin [μm] | Number of cell pairs | Proportion of similar pairs for peak amplitude (%) | Proportion of similar pairs for total signal (%) |
|---|---|---|---|
| 0 – 30 | 17 | 100 | 94.1 |
| 30 – 60 | 87 | 72.4* | 72.4 |
| 60 – 90 | 134 | 67.2** | 65.7* |
| 90 – 120 | 132 | 60.6** | 53.4** |
| 120 – 150 | 113 | 63.7** | 62.8* |
| 150 – 180 | 93 | 57** | 52.7** |
| 180 – 210 | 85 | 55.3** | 54.1** |

Asterisks indicate significant difference compared to the ratio of the first bin (Fisher's Exact Test, *$P < 0.05$, ** $P < 0.01$).

Table S2. BDNF promoter activity in response to the depolarized stimulation (KCl treatment) in different conditions

| condition | animal | number of samples | peak amplitude | total signal | slope |
|---|---|---|---|---|---|
| normal | mouse | 26 cells from 2 cultures | 7.4 ± 0.69 | 74 ± 9.4 | 1.1 ± 0.13 |
| 4 mM $Mg^{2+}$ and $Ca^{2+}$ and 100 μM PTX | rat | 25 cells from 2 cultures | 9.9 ± 0.93*** | 110 ± 11*** | 1.5 ± 0.15*** |
| normal (same as Figure 2) | rat | 123 cells from 9 cultures | 6.1 ± 0.29 | 65 ± 3.5 | 0.92 ± 0.046 |

Asterisks indicate the significant difference compared to KCl treatment in rat normal cultures (Mann–Whitney *U* test, *** $P < 0.001$).

Table S3. BDNF promoter activity in response to the patterned electrical stimulation in mouse cortical neurons.

| stimulation pattern | number of samples | total signal | | slope | | ratio of the responding cells | extent of sustained increase |
|---|---|---|---|---|---|---|---|
| 20 Hz | 35 cells from 4 cultures | before | 7.1 ± 0.89 | before | 0.14 ± 0.018 | 66 | -0.17 ± 0.12 |
| | | after | 16 ± 2.1*** | after | 0.39 ± 0.063*** | | |
| 60 Hz | 8 cells from 2 cultures | before | 7.1 ± 1.9 | before | 0.14 ± 0.038 | 25 | - |
| | | after | 7.9 ± 2.4 | after | 0.16 ± 0.040 | | |
| TB | 41 cells from 3 cultures | before | 7.1 ± 0.77 | before | 0.14 ± 0.015 | 68 | 0.12 ± 0.039 |
| | | after | 13 ± 1.6** | after | 0.23 ± 0.017*** | | |

Asterisks indicate the significant increase compared to before stimulation (Mann–Whitney *U* test, $^{**}P < 0.01$, $^{***}P < 0.001$).

Table S4. Distribution of cortical cells with similar promoter activity in response to electrical stimulation

| Bin [μm] | Number of cell pairs | Proportion of similar pairs for peak amplitude (%) | Proportion of similar pairs for total signal (%) |
|---|---|---|---|
| 0 – 30 | 5 | 100 | 80 |
| 30 – 60 | 42 | 87 | 81 |
| 60 – 90 | 60 | 74 | 75 |
| 90 – 120 | 75 | 71 | 68 |
| 120 – 150 | 81 | 75 | 72.8 |
| 150 – 180 | 56 | 63* | 62.5 |
| 180 – 210 | 53 | 63* | 60.4* |

Asterisks indicate significant difference compared to the ratio of the second bin (Fisher's Exact Test, * $P < 0.05$).

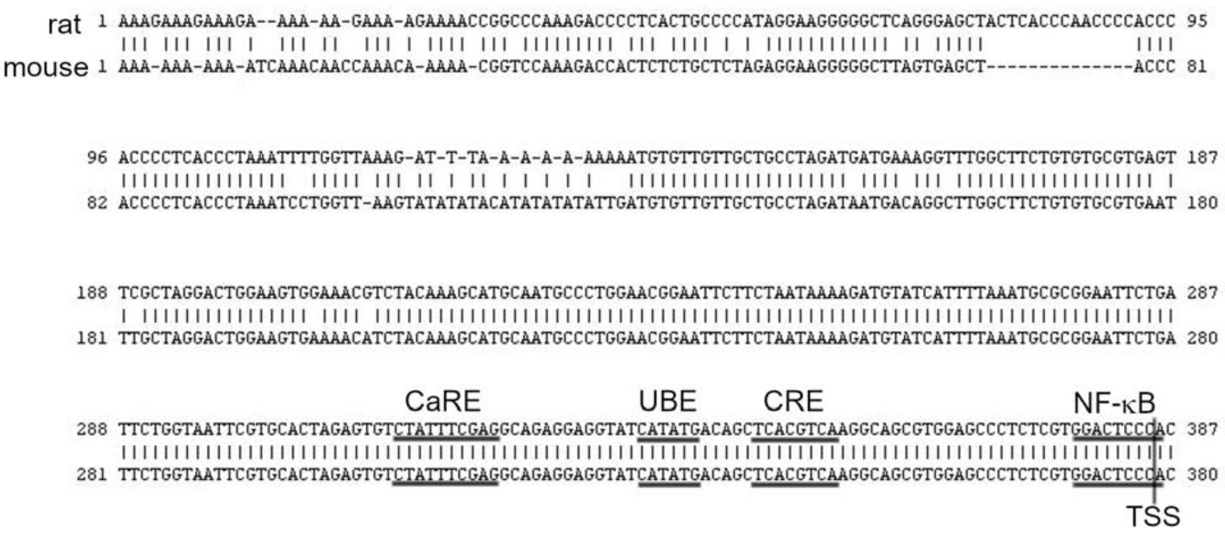


Fig. S1. Bdnf promoter IV sequence of rat and mouse. CaRE, calcium response element; UBE, upstream stimulatory factor binding element; CRE, cAMP-response element; NF-κB, nuclear factor-kappa B; TSS, transcription start site.

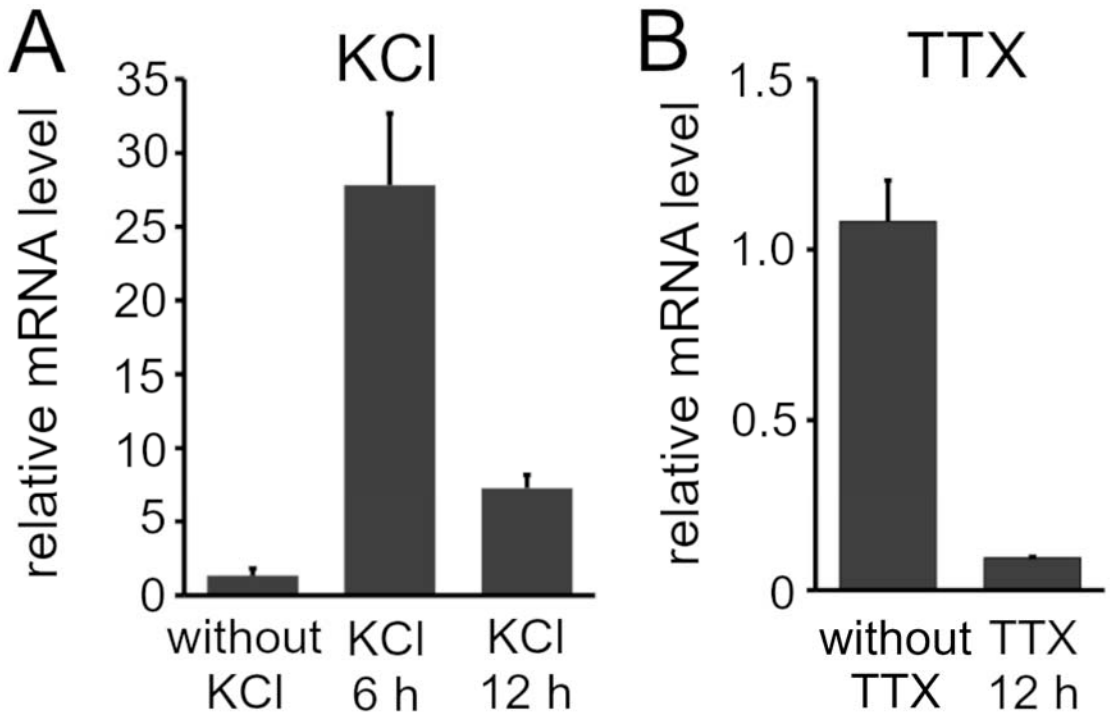


Fig. S2. Changes of endogenous Bdnf expression in response to pharmacological stimulation. mRNA abundance was examined by qRT-PCR at 1 week *in vitro* for KCl treatment (**A**) and 2 weeks *in vitro* for TTX treatment (**B**). $n = 2$ cultures for both treatments. Bars represent the mean ± SD.

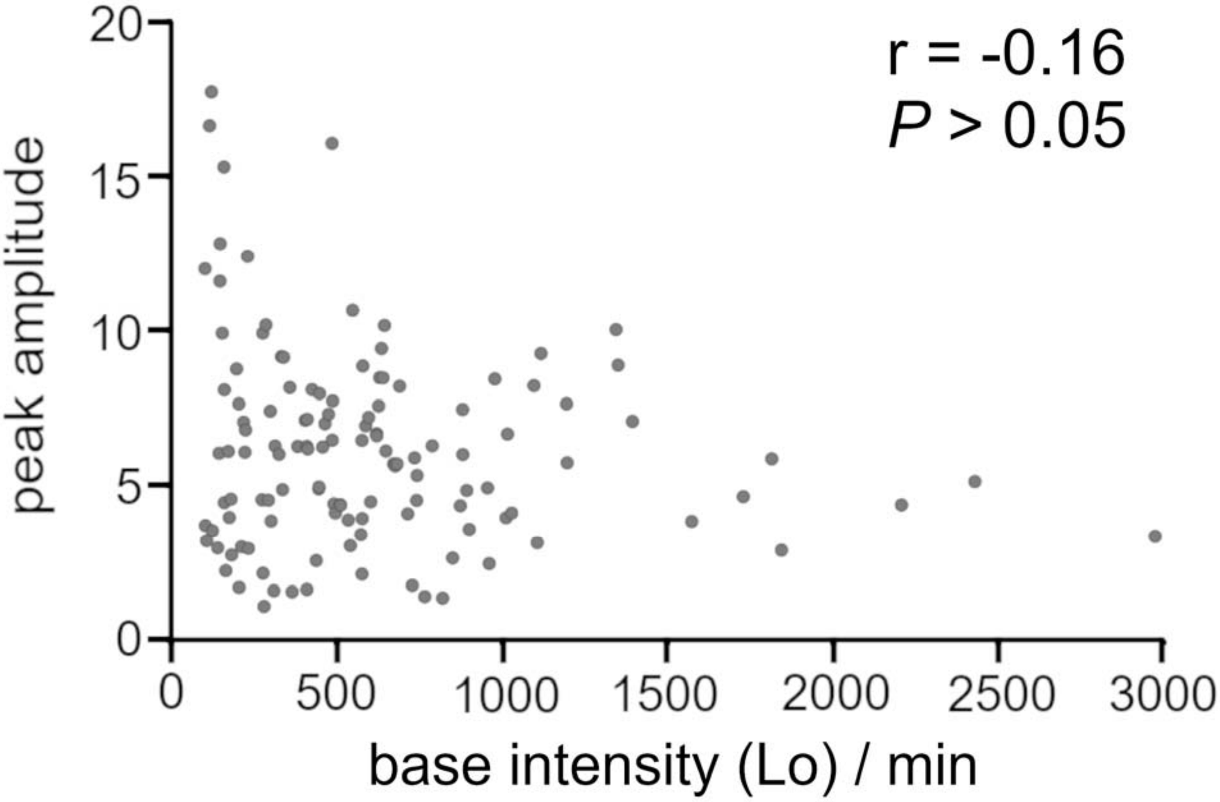


Fig. S3. Relationship between baseline intensity and peak amplitude. The baseline intensity was not related to the increased level of luciferase expression after KCl treatment. The Pearson correlation coefficient (r) is shown in the graph. The *P*-value was calculated by the test for non-correlation.

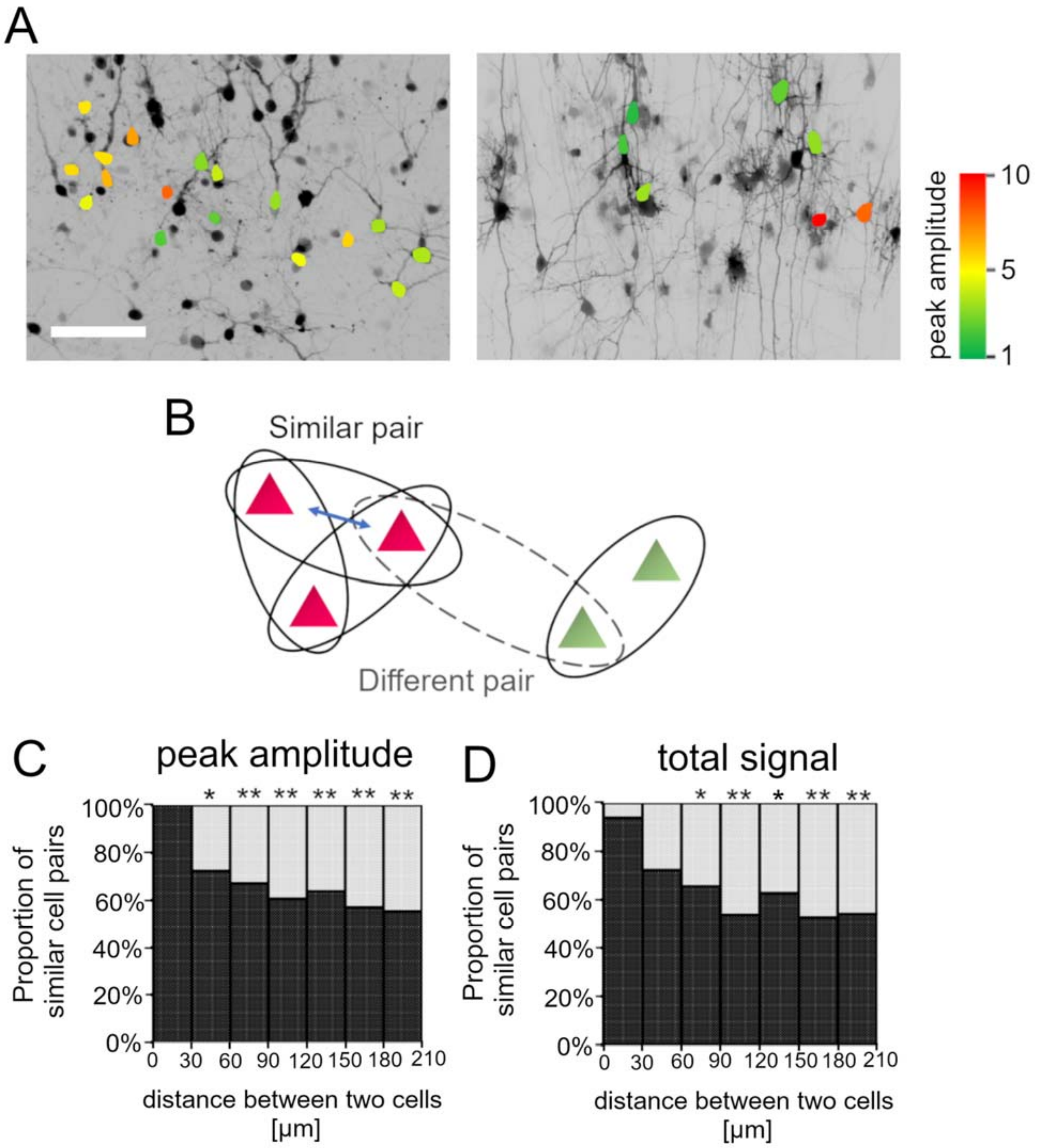


Fig. S4. Spatial distribution of cortical cells with similar promoter activity in response to KCl stimulation. **A**, Color map representations of Bdnf promoter activity in luc-positive neurons in response to KCl stimulation. Colors indicate the peak amplitudes of luc-positive cells after KCl treatment. Scale bar, 100 μm. **B**, Scheme for the analysis of spatial distributions of cortical cells with similar activity. Every cell pair was categorized as a similar pair (solid oval) or a different pair (dashed oval) according to their peak amplitudes or the total signals. The double-headed arrow indicates intercellular distance. **C, D**, The proportions of similar pairs are shown in the histogram against the intercellular distances, based on the peak amplitude (**C**) and the total signal (**D**). Asterisks indicate a significant difference compared to the proportion in the first bin (0-30 μm) (Fisher's Exact Test, * $P < 0.05$, ** $P < 0.01$).

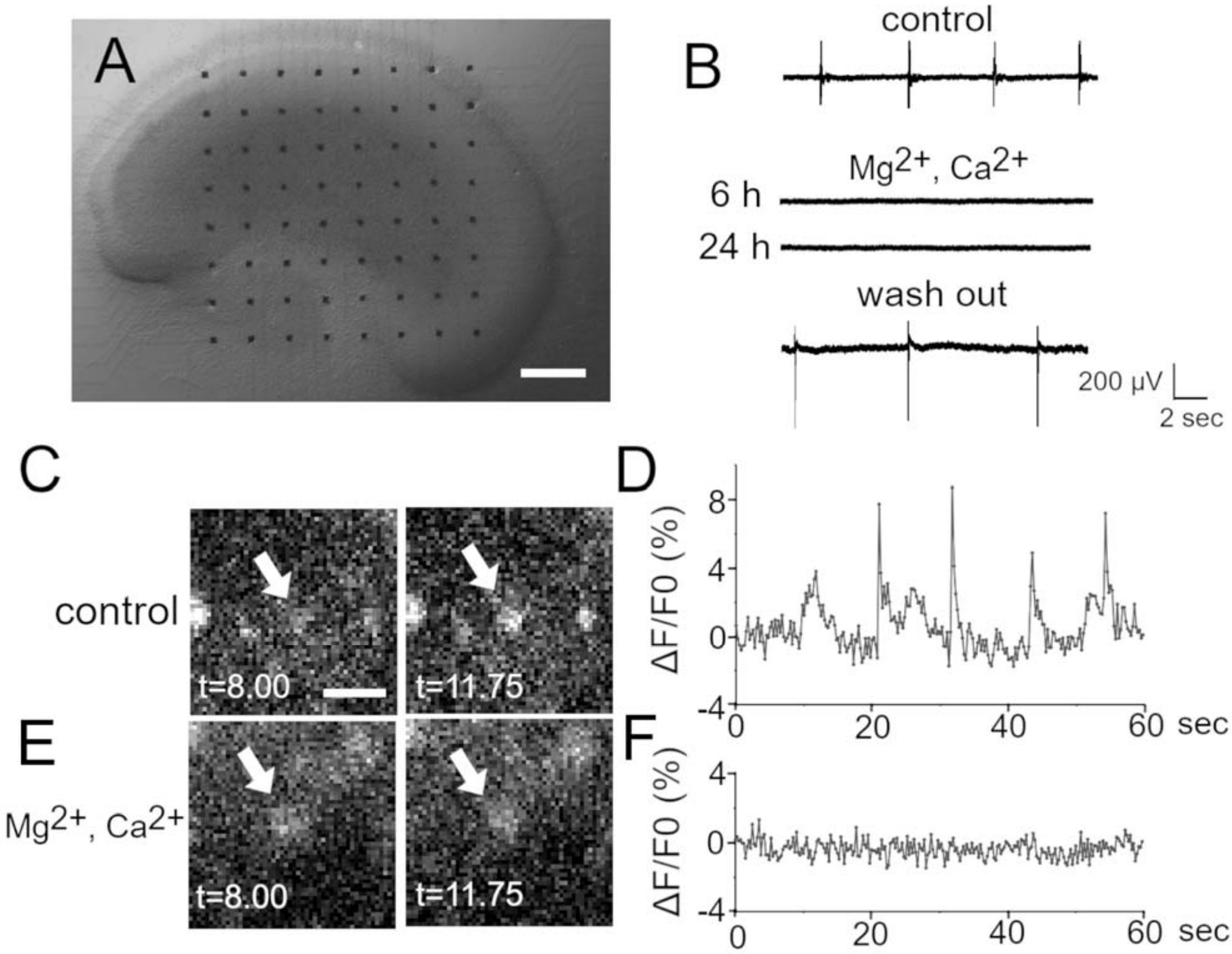


Fig. S5. Suppression of the spontaneous neuronal activity by raising the concentration of $Mg^{2+}$ and $Ca^{2+}$. **A**, Representative image of a cortical slice culture on a MED (multi electrode dish). Scale bar, 500 μm. **B**, Recording of local field potential by MED. Spontaneous activity frequently occurred around 2 weeks *in vitro* (top). The activity was almost suppressed in the presence of 4 mM $Mg^{2+}$ and $Ca^{2+}$ and 100 μM PTX for 24 h (middle), and it recovered after washing out these cations (bottom). **C-F**, Spontaneous calcium responses of OGB1-loaded neurons. Spontaneous calcium responses frequently occurred in the normal condition (**C, D**) and almost disappeared in the presence of 4 mM $Mg^{2+}$ and $Ca^{2+}$ and 100 μM PTX (**E, F**). Arrows indicate the cells analyzed for representation in the graphs. Times shown in the pictures correspond to the horizontal axes of the graphs. Scale bar, 50 μm.

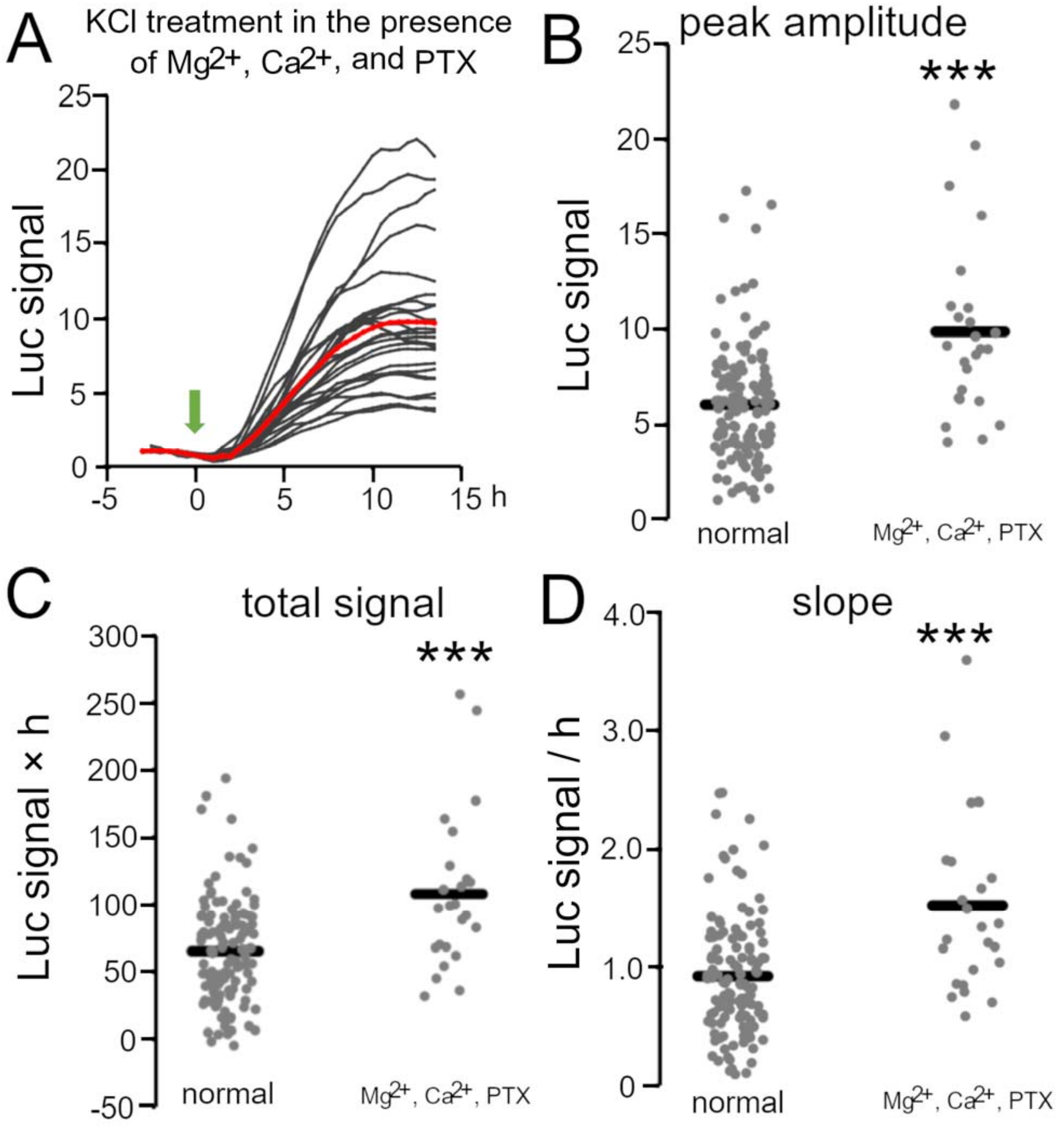


Fig. S6. BDNF promoter activity in response to depolarized stimulation in the presence of 4 mM $Mg^{2+}$ and $Ca^{2+}$ and 100 μM PTX. **A**, Time courses of luciferase signals in individual neurons with KCl treatment. Gray lines indicate luciferase signals in each neuron, and the red line shows the average. The arrow indicates the initiation of the treatment. **B-D**, Quantitative analysis of increased level for peak amplitude (B), total signal (C), and slope (D) after depolarized stimulation compared to the normal KCl treatment (see Fig. 2). The increase of promoter activity was significantly higher in the presence of 4 mM $Mg^{2+}$ and $Ca^{2+}$ and 100 μM PTX than in the normal condition, which may be due to the removal of inhibitory effects (Mann-Whitney *U* test, $***P < 0.001$). These results indicate that activity-dependent transcription was not impaired in this condition.

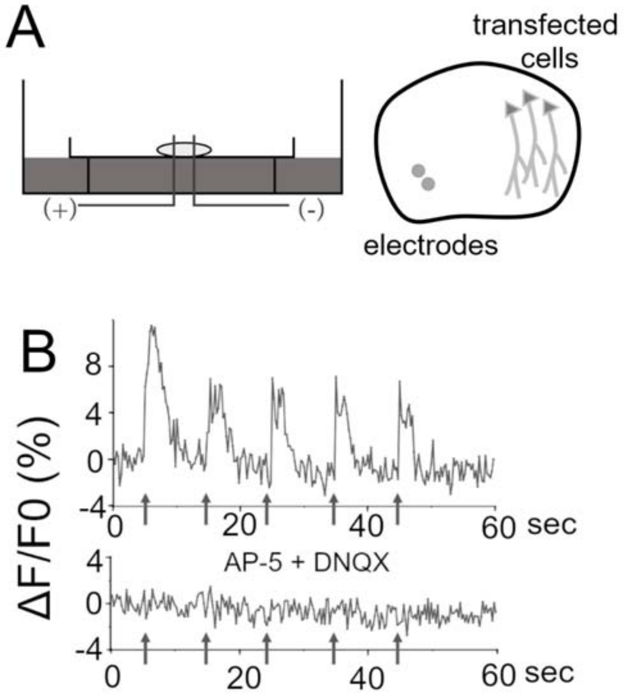


Fig. S7. Efficiency of electrical stimulations. **A**, Schematic representation of the electrode dish and arrangement of transfected neurons and electrodes. **B**, Calcium responses under electrical stimulation. Electrical stimulation (arrows) evoked calcium responses in OGB-1-loaded neurons (upper), but not in the presence of AP-5 and DNQX (lower).

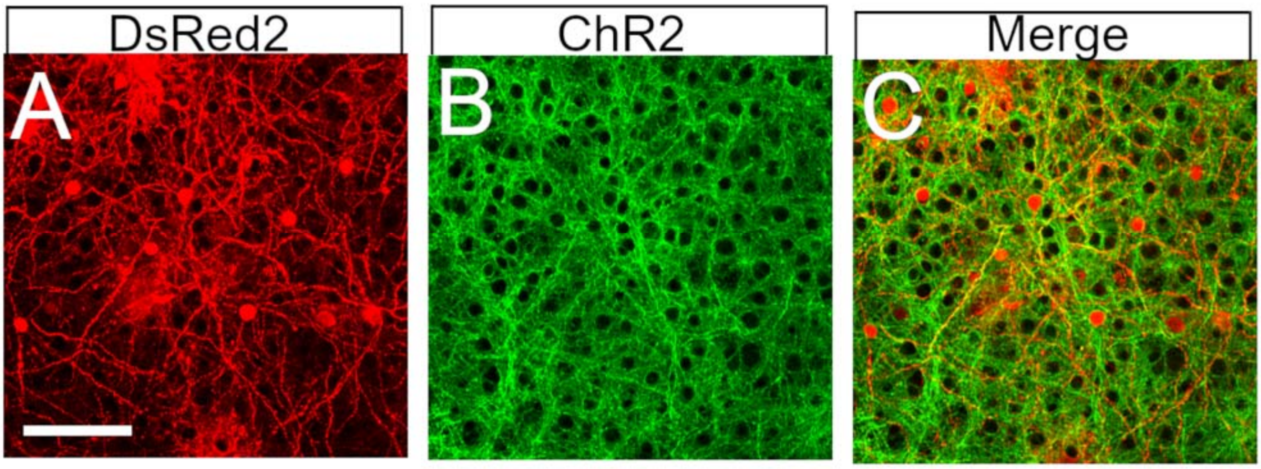


Fig. S8. Luciferase-positive neurons surrounded by ChR2 fibers in the optogenetic experiment. **A-C**, DsRed2-positive cells (**A**), ChR2-positive fibers (**B**), and merged image (**C**). Scale bar, 100 μm.

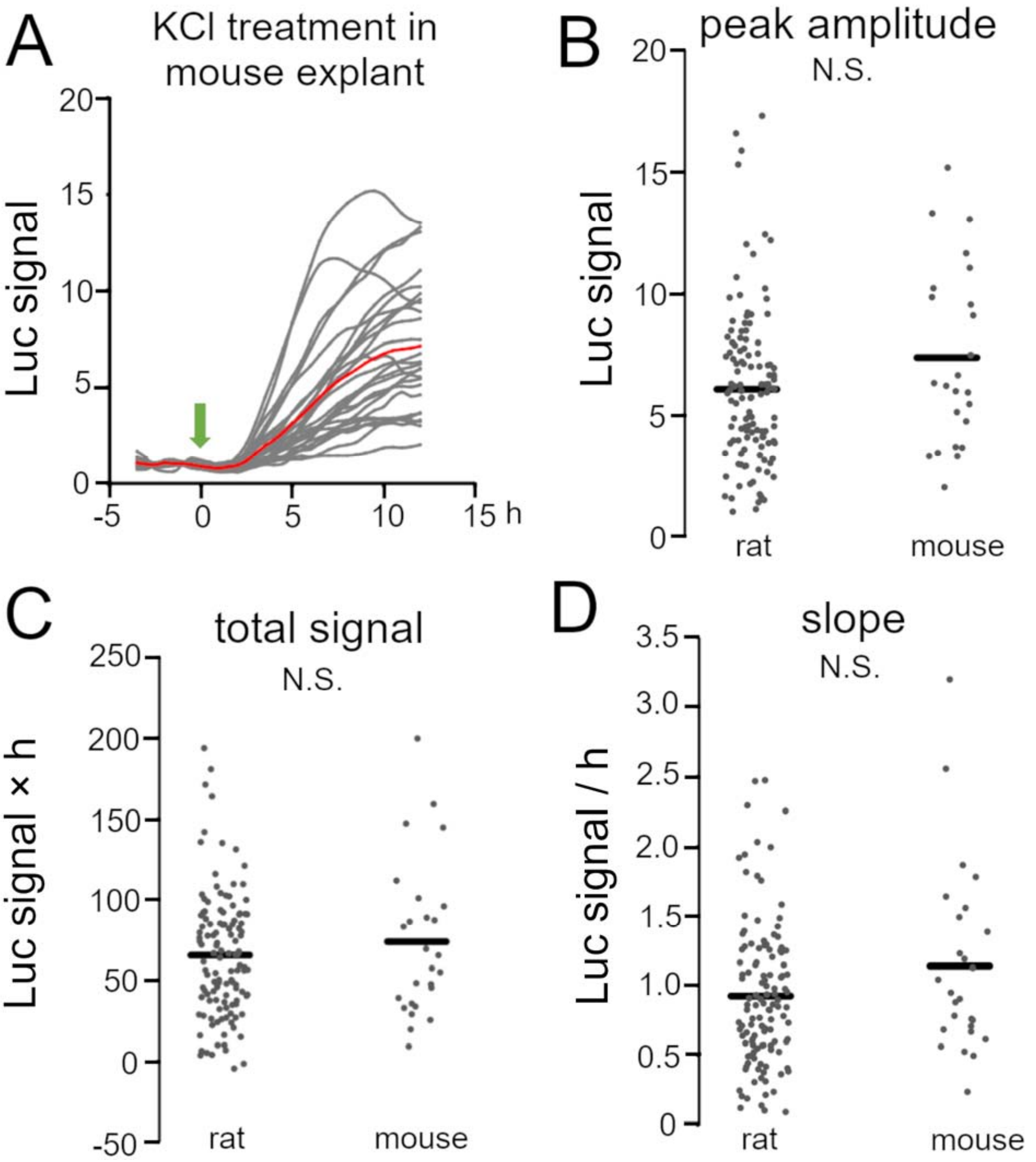


Fig. S9. Temporal properties of BDNF promoter activity in response to depolarized stimulation in mouse cortical neurons. **A**, Time courses of luciferase signals in individual mouse neurons with KCl treatment. Gray lines indicate luciferase signals in each neuron, and the red line shows the average. The arrow indicates the initiation of the treatment. **B-D**, Quantitative analysis of increased level for peak amplitude (B), total signal (C), and slope (D) after depolarized stimulation between rodents (Mann-Whitney *U* test, N.S. indicates no significance).

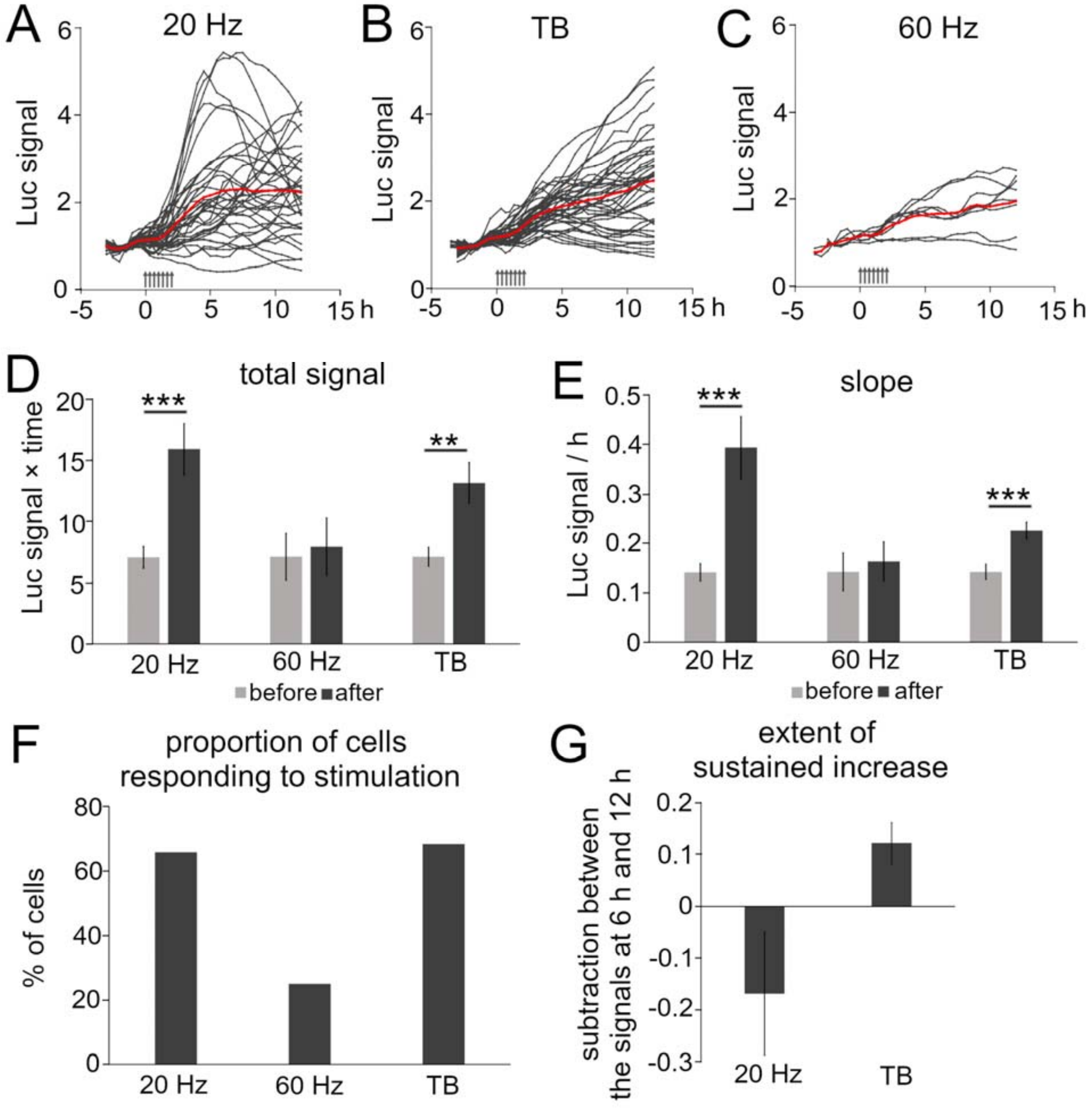


Fig. S10. BDNF promoter activity in response to patterned electrical stimulation in mouse cortical neurons. **A-C**, Time courses of luciferase signals in individual neurons for 20 Hz (**A**), 60 Hz (**B**), TB stimulation. (**C**). Gray lines indicate luciferase signals in each neuron, and the red line shows the average. **D, E**, Increased levels for total signals (**D**) and slope (**E**). **F**, Ratio of cells responding to each stimulation. **G**, Extent of the sustained increase for 20 Hz and TB stimulation. Asterisks indicate a significant difference between the signals before and after stimulation (D, E) (Mann-Whitney *U* test, $^{**}P < 0.01$, $^{***} P < 0.001$). Bars represent the mean ± SEM.

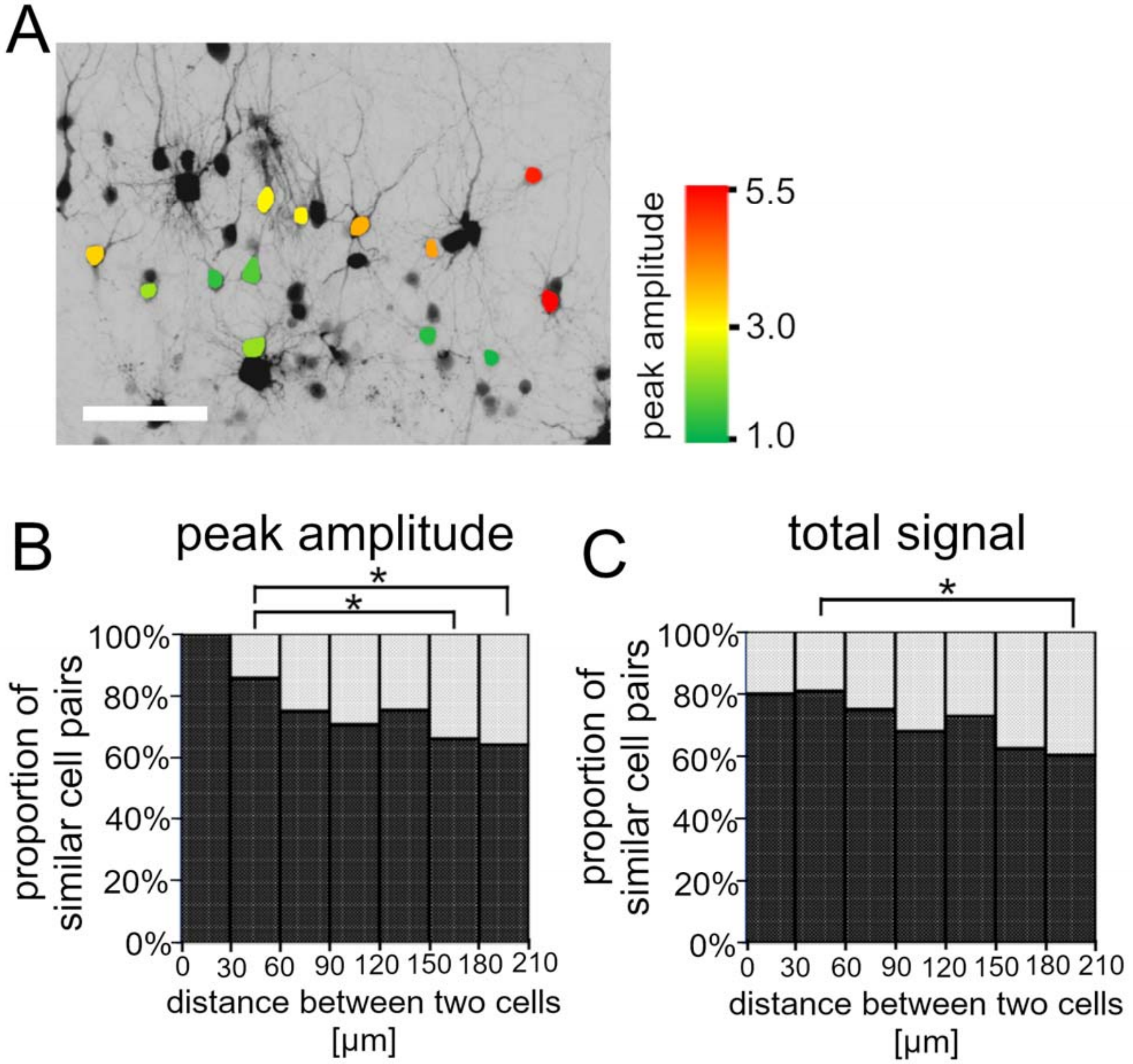


Fig. S11. Spatial distribution of cortical cells with similar promoter activity in response to patterned electrical stimulation. **A**, Color map of Bdnf promoter activity in analyzed neurons in response to the electrical stimulation. Colors indicate the peak amplitude after stimulation. Scale bar, 100 µm. **B, C**, Histograms of the proportion of similar pairs calculated from peak amplitude (**B**) and total signal (**C**). Analysis was performed from data obtained by stimulation with 20 Hz, 10 Hz, and TB. An asterisk indicates a significant difference compared to the proportion in the bin of 30-60 µm (Fisher's Exact Test, * $P < 0.05$).
